# BRC-2/BRCA2-RIPR-1 mediated constraints on homologous recombination execution are spatiotemporally regulated during meiosis

**DOI:** 10.64898/2026.06.19.733317

**Authors:** Nicola Silva, Mário Špírek, Magdaléna Ledvinová, Neeraj Bhavani Aniyan Bhavana, Jitka Blazicková, Sowmya Sivakumar Geetha, Verena Jantsch, Lumir Krejčí

## Abstract

Homologous recombination-dependent processing of meiotic double-strand breaks is crucial to generate functional gametes. The breast and ovarian cancer susceptibility gene *BRCA2* is essential for the loading of RAD51-DMC1 recombinases at DNA breaks to allow accurate repair. Here, we identify the novel protein RIPR-1 as a binding partner of *Caenorhabditis elegans* BRC-2/BRCA2. RIPR-1 and BRC-2 form an obligate complex both *in vitro* and *in vivo* and display interdependent loading in developing oocytes. Loss of *ripr-1* results in the complete destabilization of BRC-2, while *brc-2* is only partially required to preserve RIPR-1 stability. Strikingly, we found a dramatic accumulation of RAD-51 in late pachytene cells of both *ripr-1* and *brc-2* nulls, unveiling a BRCA2-independent pathway for recruitment of RAD-51 at resected meiotic DNA breaks. RIPR-1 enhances BRC-2 binding to single-stranded DNA and unlike the individual components, the assembled RIPR-1-BRC-2 complex can also bind double-stranded DNA *in vitro*. Surprisingly, we observe that RAD-51-dependent crossover designation still occurs in absence of RIPR-1-BRC-2, indicating partial, and stage-dependent constraints on homologous recombination execution. Altogether, our work identifies a novel BRC-2 interactor and sheds new light on HR-mediated regulation of DNA repair during meiosis.

## MAIN

Generation of euploid gametes depends on the accurate partitioning of the parental chromosomes into the daughter cells during meiosis. For this to be accomplished, an ensemble of tightly coordinated events unfolds during meiotic progression, culminating in the formation of physical attachments between the homologous chromosomes called crossovers (COs). COs arise from homologous recombination (HR)-mediated repair of DNA double-strand breaks (DSBs) introduced at meiotic entry by the Topoisomerase-like Spo11, as a physiological requirement to initiate recombination^1^. Spo11 induces hundreds of DSBs across the genome during germ cells development, however only a subset is repaired by HR to form COs (1-3/homologs pair across most species), whereas the vast majority is processed by non-CO repair pathways to restore genome integrity^2,3^. Therefore, absence of DSBs prevents CO formation and results in stochastic segregation of the homologs, causing formation of aneuploid gametes^4^.

COs are cytologically detectable as chiasmata, which provide the necessary tension for the correct biorientation of each homolog pair along the metaphase plate, ensuring equal chromosome segregation during the first meiotic division. Therefore, COs represent an absolute requirement for the accurate transmission of the chromosomal complement from one generation to another.

To become a substrate for the repair machinery, DSBs are first processed by the conserved MRN/X complex (Mre11-Rad50-Nbs1/Xrs1) to generate 3′single-stranded DNA overhangs that are stabilized by RPA, which successively exchanges with the recombinases RAD51 (homolog of *E. coli* RecA) and its meiotic homologue DMC1^5,6^. These in turn promote strand displacement and invasion of the homologous DNA template to allow accurate repair.

In most organisms, loading of RAD51 and DMC1 at resected DSBs is promoted by the breast and ovarian cancer susceptibility protein BRCA2/FANCD1, which physically interacts with both recombinases and allows their successful recruitment at resected DSBs^7–13^. Impaired *BRCA2* function results in reduced fertility, severe genome instability and is causative of cancer-prone syndromes such as Fanconi Anemia in humans, highlighting its pivotal function in preserving genome integrity and cellular homeostasis^14–17^.

In mammals, BRCA2 loading is promoted by PALB2/FANCN^18^ and only very recently the two novel factors BRME1 and MEILB2 have been identified in mice to directly regulate BRCA2 function in meiotic cells^19,20^. MEILB2 is essential for recruitment of BRCA2 at resected DSBs, it directly binds BRCA2 *in vitro* and the two proteins establish a functional interaction *in vivo*. Loss of MEILB2 abrogates loading of RAD51-DMC1 at meiotic DSBs in male mice, while female *Meilb2^-/-^* mutants are partially competent in forming DMC1-RAD51 foci and are sub-fertile. PALB2 knockout mice also display reduced male, but not female fertility, which may be a consequence of PALB2 interaction with BRCA1^21^. BRME1 forms a ternary complex with both MEILB2 and BRCA2, and its absence leads to complete abrogation of DSB repair, synapsis and CO formation during spermatogenesis^19,20^. Unlike BRCA2, which is fairly conserved across species, PALB2, MEILB2 and BRME1 do not show conservation of aminoacidic sequence outside vertebrate taxa, suggesting a wide diversification in the protein network assisting BRCA2 function and significantly hindering the identification of functional homologs in other species.

A functional homolog of mammalian BRCA2 was identified many years ago in *Caenorhabditis elegans* (*brc-2*) where it was shown to bind RAD-51 and to be essential for its loading, thus ensuring accurate DNA repair during meiosis^22^. However, despite its longstanding identification, no other factors involved in regulating BRC-2 function have been found in worms, leaving several aspects of its roles in the germ line largely unexplored.

In *C. elegans*, DMC1 is not present and therefore HR entirely relies on RAD-51, which shows a reproducible expression pattern in the germ line forming chromatin associated foci at meiotic entry (Transition Zone, TZ, corresponding to Leptotene-Zygotene stages) that peak in number by early-mid Pachytene (EP-MP respectively), to then disengage from DNA by late Pachytene (LP), indicating successful completion of repair^23,24^.

Chromosomes become increasingly condensed through Pachytene and Diplotene, appearing as six individual bodies by 4’,6-diamidin-2-fenylindol (DAPI) staining in Diakinesis (the last stage of meiotic Prophase I preceding Metaphase I) each representing a couple of homologous chromosomes held together by a chiasma. Impaired DSB formation or CO-mediated pathway result in the formation of twelve achiasmatic univalents with a regular morphology^4,25,26^, whereas dysfunctional DSB processing causes dramatic abnormalities in chromosome morphology and number, which can appear as clumped chromatin fusions and/or chromosome fragmentation, as observed in *brc-2* and *rad-51* mutants^22,27^.

We have previously generated a fully functional *FLAG::brc-2* tagged line by CRISPR/Cas9^28^, that we have exploited to identify putative BRC-2 interactors in *C. elegans via* a biochemical approach. Here, we report the identification of the uncharacterized open reading frame *D1007.8* as a BRC-2 interactor, which we renamed *ripr-1* (***r***ecombination ***i***ntermediates ***<u>pr</u>***ocessing-***<u>1</u>***). Removal of *ripr-1* triggers nearly complete embryonic lethality due to formation of highly aberrant chromosome structures in Diakinesis nuclei. Removal of SPO-11 suppresses aberrant chromatin figures observed in *ripr-1* mutants, indicating that meiotic DSBs are formed but not processed correctly. We found that RIPR-1 establishes an obligate complex with BRC-2 and they display a largely overlapping and mutually dependent localization in the germ line. Loss of RIPR-1 impairs RAD-51 loading at meiotic entry, however cells at late pachytene stage display a sudden and extensive accumulation of RAD-51, a similar and unexpected phenotype that we also observe in *brc-2* null mutants. This indicates that BRC-2-RIPR-1 activity is not required, *per se*, to promote global recruitment of RAD-51 to DSBs and rather hints at stage-specific regulation of RAD-51 loading dynamics exerted by the RIPR-1–BRC-2 complex. Surprisingly, HR-dependent recruitment of pro-CO factors is only mildly reduced in *ripr-1* and *brc-2* mutants and finally, we show that blocking binding of BRC-2 to RAD-51 does not impair loading or stability of RIPR-1–BRC-2, suggesting that assembly of the complex does not require its pro-recombinogenic functions. Altogether, our work identifies a novel interactor of BRC-2 in *C. elegans* and it crucially highlights previously unanticipated functions of BRC-2/BRCA2 in regulating meiotic DNA repair.

## RESULTS

### RIPR-1 is a novel factor essential for fertility

In *C. elegans*, very little is known about the BRC-2 interactome. To identify potential interactors, we used our functional *FLAG::brc-2* strain to perform immunoprecipitation experiments followed by mass spectrometry. Among the most prominent hits (Supplementary Table 1), we identified the uncharacterized ORF *D1007.8*, which we named *ripr-1* (***r****ecombination **i**ntermediates **<u>pr</u>**ocessing-**<u>1</u>***). The *ripr-1 gene* encodes for a small protein of approximately 30 kD. *In silico* analysis using AlphaFold identified the presence of a putative oligonucleotide/oligosaccharide-binding fold domain (OB-fold) within the N-terminal half of RIPR-1 (Fig. 1A), a structural motif found in DNA-binding and DNA repair proteins, including BRC-2 and RPA-1^29,30^. This prompted us to further pursue RIPR-1 characterization.

**Figure 1.**
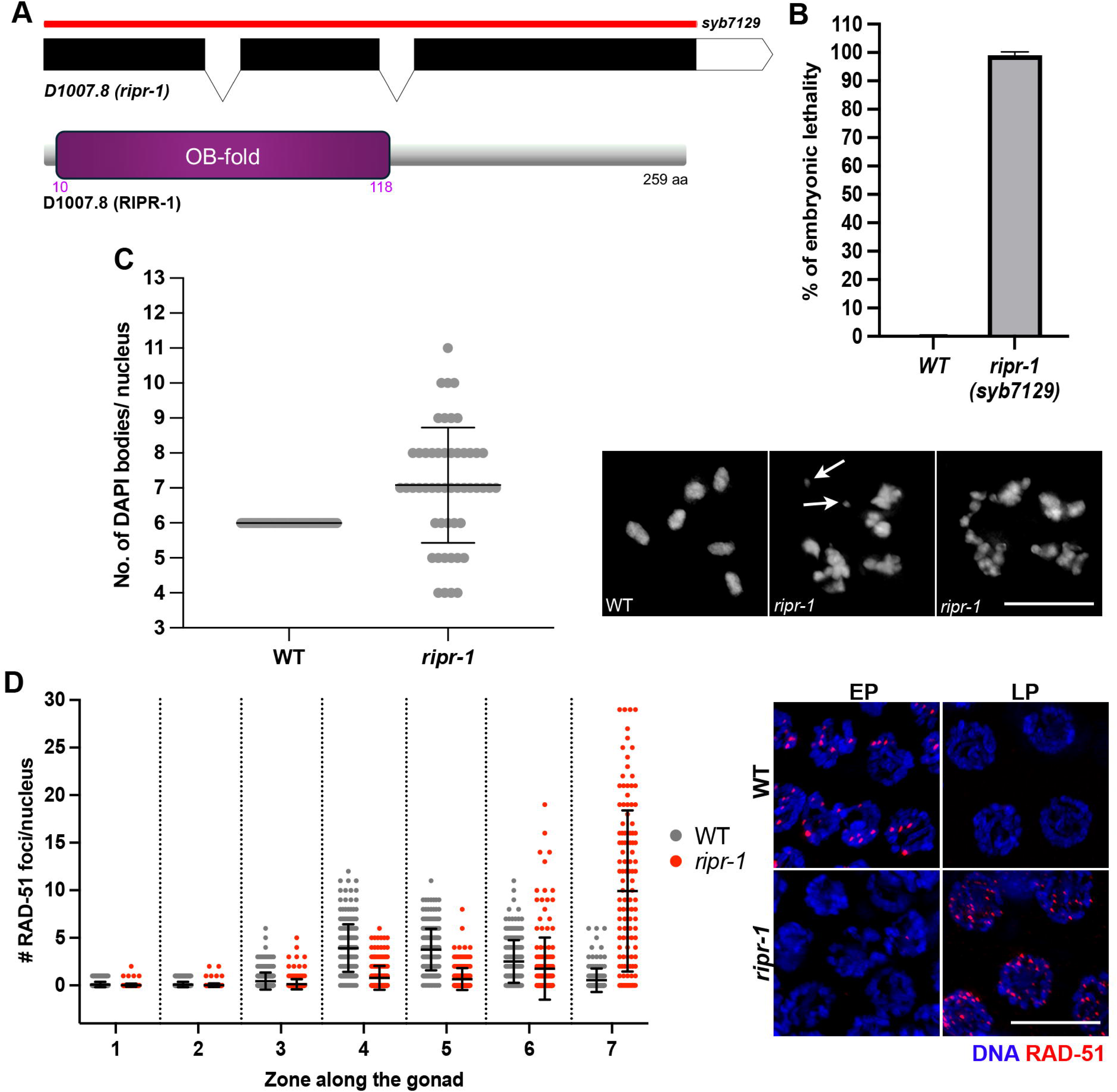
RIPR-1 is a novel factor essential for fertility. **(A)** Schematic representation of *ripr-1*(*D1007.8*) locus and protein domain organization. The *ripr-1(syb7129)* carries a full deletion of the *D1007.8* locus as depicted by the red line. Purple numbers indicate aminoacidic residues spanning the OB-fold domain. **(B)** Quantification of embryonic lethality in *ripr-1* mutants and WT controls. Bar reports average with S.D. **(C)** Left: quantification of DAPI bodies in Diakinesis nuclei of the indicated genetic backgrounds. Bars indicate average and S.D. Right: representative images of DAPI-stained Diakinesis nuclei in *ripr-1* mutants and WT controls. Arrows indicate chromosome fragments. Scale bar 5 μm. **(D)** Left: quantification of RAD-51 foci in WT controls and *ripr-1* mutants. Chart shows the number of RAD-51 foci/nucleus in the indicated regions of the gonad and bars indicate average with S.D. Right: representative images of nuclei from the indicated stages (EP= early pachytene, LP= late pachytene) and genetic backgrounds stained for RAD-51 (red) and counterstained by DAPI (blue). Scale bar 5 μm.

Since no mutant alleles were available, we generated the complete *ripr-1(syb7129)* knockout (Fig. 1A) and assessed viability levels. Compared to WT animals, *ripr-1* null mutants produced roughly normal broods on average (*ripr-1*: 194.8±16.4 SD. – WT: 259.4±55.1 SD., *p=0.05*, not significant). However, only 0.9% of the laid embryos hatched (13/1364) (Fig. 1B) and none of these larvae reached adulthood, indicating that RIPR-1 plays essential functions.

To examine the meiotic defects underlying this phenotype, we interrogated CHK-2-mediated phosphorylation of SUN-1 at Serine 8 (pSUN-1^S8^), which marks a checkpoint mechanism monitoring the timely execution of meiotic tasks during meiotic Prophase I^31^. Under dysfunctional DSB induction/repair or synapsis, CHK-2 activity, which is normally shut down by EP, is substantially prolonged, resulting in the extended phosphorylation of SUN-1^S8^ until LP stage. Therefore, immunolabeling of pSUN-1^S8^ can be used as a sensitive readout for the completion of early meiotic events. We found that SUN-1^S8^ phosphorylation persisted throughout MP stage in the germ line of *ripr-1* mutants compared to WT animals (86.8%±5.02 S.D. vs. 57.9%±2.58 S.D. respectively, *p=0.016\**), suggesting delayed meiotic progression (Supp. Fig. 1A). To identify the defect(s) underlying pSUN-1^S8^ extension, we assessed Synaptonemal Complex (SC) assembly by cytological analysis of the axial component HTP-3 and the central element SYP-1. The SC is a proteinaceous structure that keeps the homologous chromosomes in proximity and allows CO formation. It consists of lateral (HTP-3, HIM-3, HTP-1/-2 and cohesins)^4,32–37^ and central (SYP-1,-2, −3, −4, −5, −6 and SKR-1/-2)^24,38–42^ components, that are assembled progressively along paired homologous chromosomes, with the axial components loaded prior to the central elements. Regions decorated by axis markers but devoid of central elements indicate absence of synapsis. To quantify the extent of synapsis, gonads were divided into six equal zones from TZ to Diplotene entry, and synapsis was assessed in all nuclei in each zone. Nuclei displaying fully overlapping HTP-3/SYP-1 signals were scored as fully synapsed, while nuclei displaying zones positive for HTP-3 but lacking SYP-1 were quantified as partially synapsed^43^.

While substantial levels of synapsis were achieved in *ripr-1* mutants, nuclei exhibiting incomplete synapsis were frequently scattered throughout the germ line (Supp. Fig. 1B). In particular, we observed that this phenotype was exacerbated in zone 6 (LP), where nearly 90% of nuclei displayed incomplete synapsis in the *ripr-1* mutants. Previous work has shown that reduced, but not abrogated establishment of COs triggers precocious removal of the SC along the chromosomes that failed undergoing recombination in LP^44,45^, and therefore the high levels of incomplete synapsis observed in *ripr-1* mutants may suggest possible defects in the efficient establishment of COs.

Consistent with the high levels of embryonic lethality and the incomplete synapsis, DAPI staining showed that Diakinesis chromosomes in *ripr-1* mutants appeared mostly as unstructured chromatin masses, fragments, or univalents (Fig. 1C), indicating that meiotic DSBs are likely formed but not properly processed^22,46–48^. Since *C. elegans* lacks a direct cytological marker for DSBs, RAD-51 is commonly used as an indirect proxy to monitor formation and resolution of recombination intermediates^23,24,49^. We monitored RAD-51 foci by dividing the gonads in seven equal zones spanning the pre-meiotic compartment through Diplotene entry and counting the number of foci/nucleus (Supp. Fig. 2A). Loading of RAD-51 was severely reduced from TZ through MP (Fig. 1D zones 1-5 and Supp. Fig. 2A), followed by a sudden and dramatic accumulation of foci at MP-LP, extending into Diplotene (Fig. 1D zones 6-7 and Supp. Fig. 2A). This zone overlaps with execution of apoptotic cell death, which under normal conditions of growth culls meiocytes most likely as a control system to maintain homeostasis of the gonadal tissue^50^. However, defects in chromosome synapsis^51^ and/or DNA repair^52^ can greatly increase apoptosis levels. DNA damage triggers activation of the pro-apoptotic pathway through CEP-1, the only worm ortholog of mammalian *p53*, which is required for apoptosis execution^52^. To assess whether the extensive DNA damage observed in *ripr-1* mutants depended on apoptosis, we generated the *ripr-1 cep-1* double mutants and quantified RAD-51 foci formation. We observed extensive RAD-51 foci formation in the *ripr-1 cep-1* doubles (Supp. Fig. 2B) similar to the *ripr-1* single mutants, indicating that persistence of recombination intermediates does not result from damage-induced apoptosis. Collectively, these findings show that RIPR-1 is a novel putative BRC-2 interactor whose activity is essential to maintain fertility and to promote timely loading of RAD-51, thereby facilitating proper processing of meiotic recombination intermediates.

**Figure 2.**
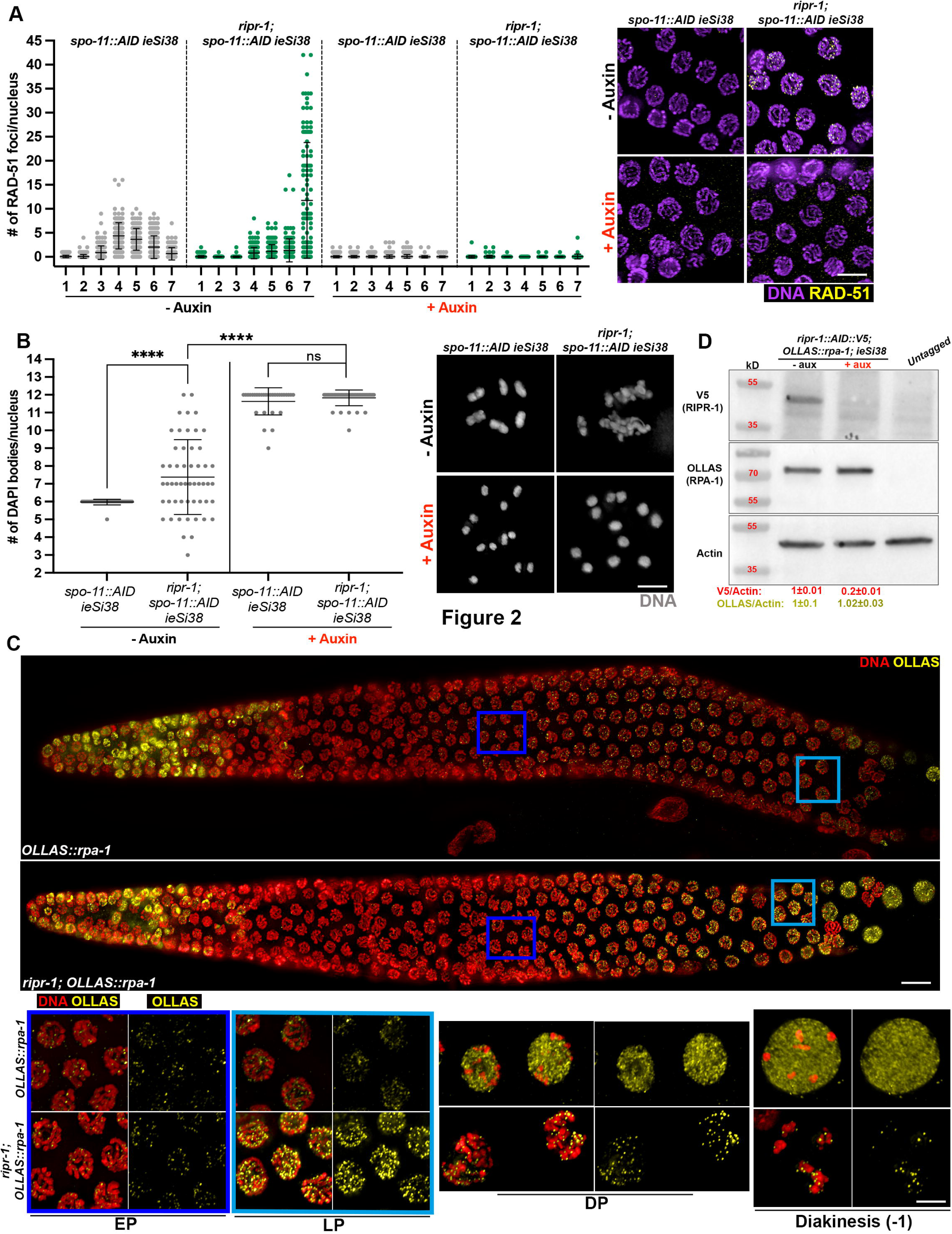
Loss of RIPR-1 results in impaired processing of SPO-11-dependent DSBs. **(A)** Left: quantification of RAD-51 foci in the indicated genetic backgrounds and exposure to auxin. Chart shows the number of RAD-51 foci/nucleus in the indicated regions of the gonad (X axis) and bars indicate average with S.D. Right: representative images of nuclei from mid/late-pachytene stage in the indicated genetic backgrounds and auxin exposure conditions stained for RAD-51 (yellow) and counterstained by DAPI (purple). Scale bar 5 μm. **(B)** Left: quantification of DAPI bodies in Diakinesis nuclei of the indicated genetic backgrounds and exposure conditions to auxin. Bars indicate average and S.D. Asterisks denote statistical significance assessed by T test (*\*\*\*\*p<0.0001, ns*= not significant). Right: representative images of DAPI-stained Diakinesis nuclei in *ripr-1* mutants and WT controls before and after auxin-induced degradation of SPO-11::AID. Scale bar 5 μm. **(C)** Whole-mount gonads of the indicated genotypes stained for OLLAS::RPA-1 (yellow) and counterstained by DAPI (red). Scale bar 5 μm. Colored squares indicate regions magnified below for EP (Early Pachytene) and LP (Late Pachytene), followed by magnified representative images of Diplotene (DP) and Diakinesis nuclei. **(D)** Western Blot on whole cell extracts from *ripr-1::AID::V5; OLLAS::rpa-1; ieSi38* animals before and after 24h exposure to auxin, and untagged negative controls. The membrane was probed with the indicated antibodies, and the ration of V5/Actin (red) and OLLAS/Actin (dark yellow) is reported below, ±S.D. The analysis was performed in biological duplicates and shows no significant variation of total RPA-1 protein levels upon loss of RIPR-1.

### Loss of RIPR-1 impairs processing of SPO-11-dependent DSBs and causes accumulation of unrepaired ssDNA

Given the perturbed loading of RAD-51, we next wondered whether this accumulation occurred at meiotic DSBs or following DNA damage ensuing during mitotic replication. To this end, we crossed the *ripr-1* mutant into the *spo-11::AID ieSi38* degron strain that carries TIR1 expression driven by *sun-1* regulatory elements (*ieSi38*), in which exposure to auxin triggers efficient degradation of AID-tagged SPO-11 thereby abrogating meiotic break formation^53,54^. RAD-51 foci were largely abolished in *ripr-1; spo-11::AID ieSi38* animals exposed to auxin for 24h (Fig. 2A) and Diakinesis nuclei of *ripr-1* mutants were indistinguishable from *spo-11::AID* worms (Fig. 2B), indicating that aberrant RAD-51 accumulation and the consequential formation of chromosome abnormalities are due to impaired processing of meiotic DSBs.

Processing of SPO-11-dependent breaks generates 3′-single stranded DNA ends through nucleolytic resection, providing a substrate for RPA-1 binding and stabilization. Subsequently, RPA-1 is displaced by RAD-51, which forms filaments on the ssDNA to promote homology search and strand exchange with the homologous donor template.

To address whether loss of RAD-51 accumulation at early meiotic stages in *ripr-1* mutants was due to impaired resection, we monitored the localization of two components of the MRN/X complex, MRE-11^46,55^ and RAD-50^48,56^, using previously established functional CRISPR-tagged lines^28,57^. We detected no gross alterations in the loading/expression of MRE-11::GFP (Supp. Fig. 3A) or RAD-50::FLAG (Supp. Fig. 3B), suggesting that the MRN/X complex localizes with meiotic chromatin independently of RIPR-1.

**Figure 3.**
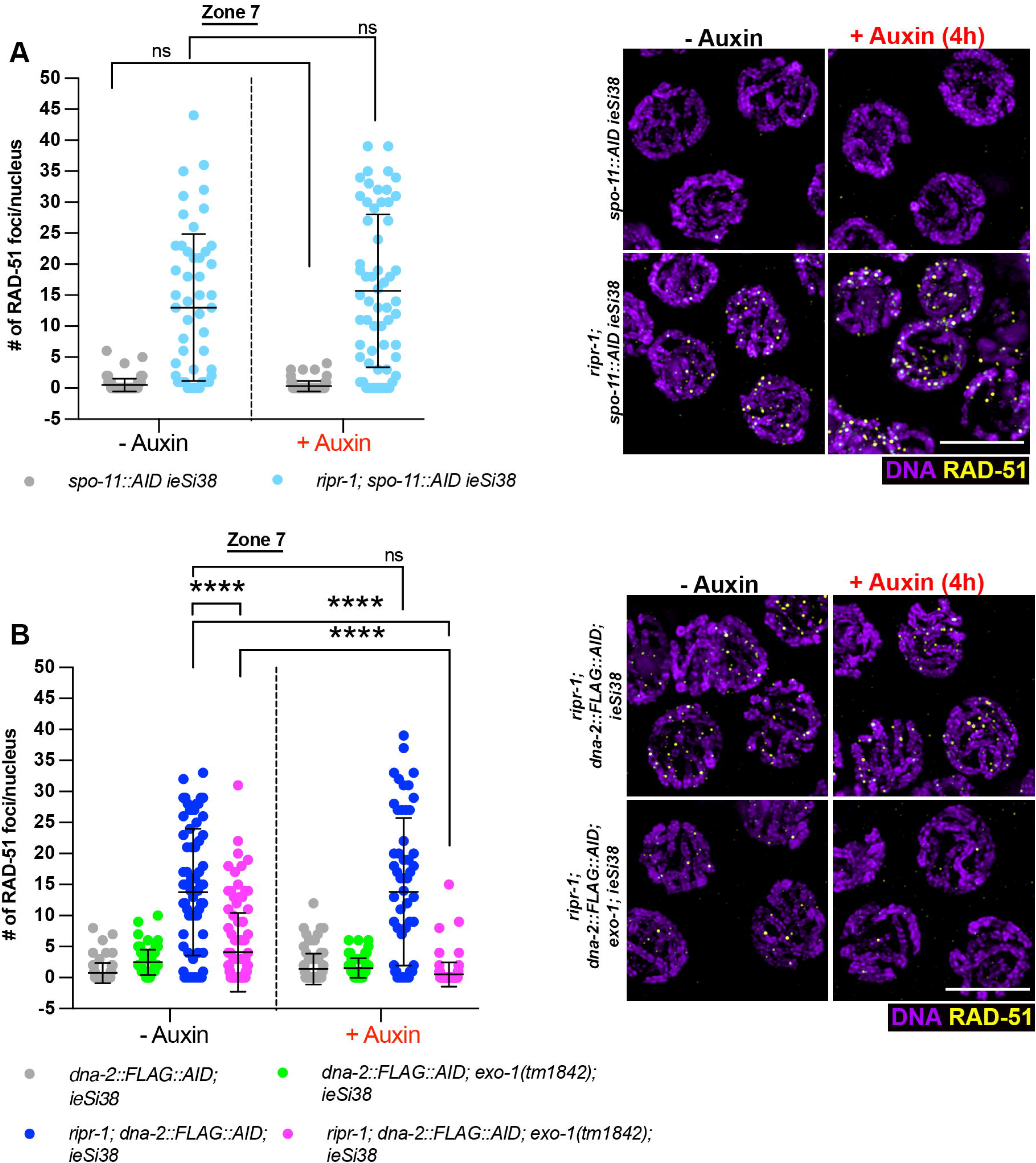
Late RAD-51 accumulation in *ripr-1* mutants depends on long range resection. **(A)** Left: quantification of RAD-51 foci number in zone 7 (corresponding to LP) in *ripr-1* mutants before and after short depletion of SPO-11 (4h). Bars depict average with S.D., “*ns*” indicates not significant differences, as assessed by T test. Right: representative images of late pachytene nuclei from the indicated genetic background and exposure conditions to auxin, immunoassayed for RAD-51 (yellow) and counterstained by DAPI (purple). Scale bar 5 μm. **(B)** Left: quantification of RAD-51 foci number in zone 7 (corresponding to LP) in the indicated genetic background and exposure conditions to auxin. Bars indicate average with S.D., “*ns*” indicates not significant differences, and asterisks denote statistical significance as assessed by T test (*\*\*\*\*p<0.0001*). Right: representative images of late pachytene nuclei from the indicated genetic background and exposure conditions to auxin, immunoassayed for RAD-51 (yellow) and counterstained by DAPI (purple). Scale bar 5 μm.

Consistent with functional end-resection by MRN/X complex, we next monitored OLLAS::RPA-1^58^ recruitment as an indirect readout of ssDNA formation. Initial loading of OLLAS::RPA-1 was not perturbed in *ripr-1 mutants* (Fig. 2C, EP), corroborating that the defective RAD-51 foci formation is not due to impaired generation of ssDNA. Strikingly, while in control animals OLLAS::RPA-1 progressively disengaged from chromatin during MP/LP and redistributed into the nucleoplasm (Fig. 2C, LP and DP), in *ripr-1* mutants it dramatically accumulated as discrete chromatin-associated foci that persisted until Diakinesis (Fig. 2C). This indicates that ssDNA is generated and stabilized by RPA-1 but its subsequent exchange with RAD-51 is prevented, resulting in the extensive accumulation of unrepaired ssDNA.

Next, we generated the functional *ripr-1::AID::V5* degron allele, which enabled efficient auxin-induced depletion of the RIPR-1 protein. Western blot analysis confirmed successful depletion of RIPR-1 and showed that the total OLLAS::RPA-1 protein pool remained largely unchanged upon removal of RIPR-1 (Fig. 2D), indicating that extensive RPA-1 accumulation observed cytologically reflects a global re-localization to ssDNA rather than increased protein levels.

Previous work from our lab has shown that in the *C. elegans* germ line, SPO-11-dependent DSBs are generated both at early and later stages, with the latter serving as a substrate for long-range resection (LRR) driven by the partially redundant DNA-2 and EXO-1-mediated activities^54^. In *ripr-1* mutants, we observed extensive RAD-51 recruitment only at MP/LP stage (Fig. 1 and Supp. Fig. 2), whereas OLLAS::RPA-1 displayed increasing levels throughout the germ line (Fig. 2), which were concomitantly observed with RAD-51 foci in LP cells (Supp. Fig. 4A), indicating deeply perturbed HR kinetics. This pattern suggests that i) *de novo* formation of meiotic DSBs, and thus newly generated ssDNA, could be taking place in MP/LP, where the constraints imposed by RIPR-1 on RAD-51 loading may no longer be active, ii) or that end resection could proceed continuously due to inability to load RAD-51 at stabilized ssDNA tails at earlier but not later stages.

**Figure 4.**
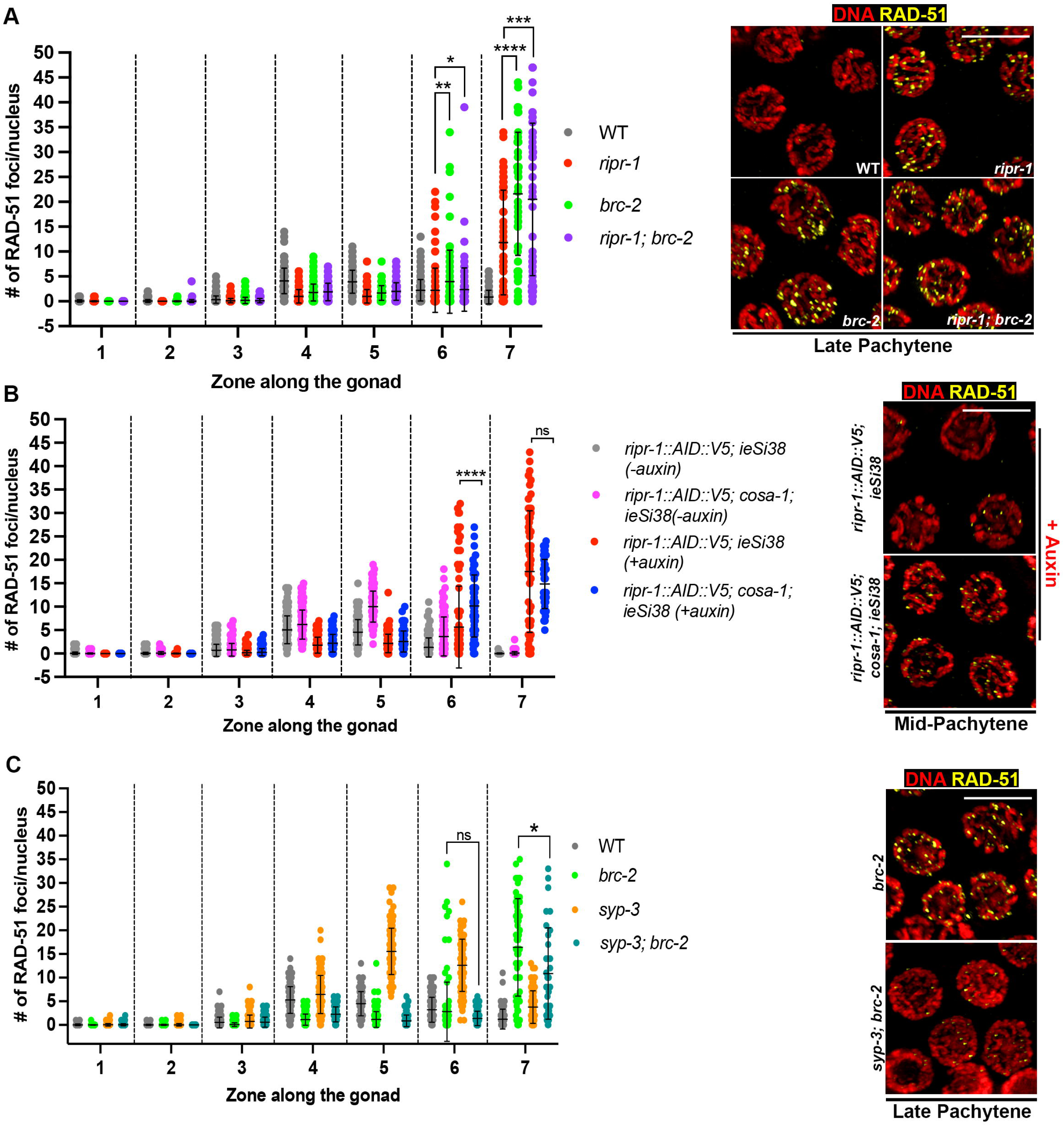
RIPR-1-BRC-2 act in the same genetic pathway. **(A)** Left: quantification of RAD-51 foci across the germ line in the indicated genetic backgrounds. Bars report average with S.D. and asterisks denote statistical significance as assessed by T test (*\*p=0.04,**p=0.0015,****p<0.0001*). Right: representative images of late pachytene nuclei from the indicated genetic background immunoassayed for RAD-51 (yellow) and counterstained by DAPI (red). Scale bar 5 μm. **(B)** Left: quantification of RAD-51 foci across the germ line in the indicated genetic backgrounds and exposure conditions to auxin. Bars report average with S.D. and asterisks denote statistical significance as assessed by T test (*\*\*\*\*p<0.0001, ns=* not significant). Right: representative images of mid-pachytene nuclei from the indicated genetic background immunoassayed for RAD-51 (yellow) and counterstained by DAPI (red). Scale bar 5 μm. **(C)** Left: quantification of RAD-51 foci across the germ line in the indicated genetic backgrounds. Bars report average with S.D. and asterisks denote statistical significance as assessed by T test (*\*p=0.01, ns=*not significant). Right: representative images of late pachytene nuclei from the indicated genetic background immunoassayed for RAD-51 (yellow) and counterstained by DAPI (red). Scale bar 5 μm.

To test these hypotheses, we first performed a short (4h) SPO-11 depletion in the *ripr-1; spo-11::AID ieSi38* worms and analysed RAD-51 localization in LP nuclei (zone 7). This brief exposure time to auxin ensures that the depletion coincides with the stage analysed^54^. Depletion of SPO-11 did not reduce RAD-51 foci in *ripr-1* worms, ruling out the possibility that these foci originated from newly induced DSBs in LP (Fig. 3A).

As a second approach, we generated a strain in which both DNA-2 and EXO-1 activities were abrogated by combining a *dna-2::AID::FLAG* allele^54^ with the *exo-1(tm1842)* deletion mutant. These worms were exposed to auxin for 4h, sufficient to deplete DNA-2::AID::FLAG in LP cells^54^. Quantification of foci number in *ripr-1; dna-2::FLAG::AID; exo-1(tm1842); ieSi38* animals showed that strikingly, RAD-51 accumulation in LP nuclei was nearly abrogated (Fig. 3B, magenta, -auxin vs. +auxin,), indicating that late recombination intermediates in *ripr-1* mutants require LRR to load RAD-51. Interestingly, we noticed that in the *ripr-1; dna-2::FLAG::AID; exo-1(tm1842)* worms not exposed to auxin (thus corresponding to *ripr-1; exo-1* double mutants), RAD-51 foci were already significantly reduced compared to *ripr-1* single mutants (Fig. 3B, magenta vs. blueberry, -auxin), a phenotype that was not observed when only DNA-2 was removed (Fig. 3B, blueberry, + auxin vs blueberry, -auxin). These results indicate that RIPR-independent loading of RAD-51 observed in LP cells relies on LRR and highlight a predominant role of EXO-1 over DNA-2 in this process.

### ripr-1 and brc-2 act in the same genetic pathway

Our phenotypical analysis so far shows striking similarities between the *ripr-1* and *brc-2* mutants, since embryonic lethality, Diakinesis phenotype, and RPA-1 accumulation were all reminiscent of *brc-2(tm1086)* mutants^22^. To explore possible epistatic relationships between *ripr-1* and *brc-2*, we first considered the nature of the existing *brc-2(tm1086)* mutant, which is a partial deletion allele. While previous analyses suggest that *brc-2(tm1086)* likely represents a null background, we decided to generate the complete *brc-2(syb7480)* knock-out by CRISPR-Cas9, which we employed for most experiments unless otherwise indicated. The *brc-2(syb7480)* mutants recapitulated the embryonic lethality observed in the *brc-2(tm1086)* worms, at levels comparable to *ripr-1* mutants (Supp. Fig. 4B). We noticed that the average brood size in our allele was higher (*brc-2(syb7480)*: 193.4±29 S.D. – *brc-2(tm10860)*: 41.1±10 S.D.) and these worms appeared generally healthier, as the *brc-2(tm1086)* animals often display an egg-laying defective phenotype (*Egl*, eggs accumulate in the uterus and are not expelled) that we did not observe in the *brc-2(syb7480)* animals.

Next, we generated the *ripr-1; brc-2(syb7480)* double mutant and analyzed RAD-51 localization. Similar to *ripr-1* single mutants, *brc-2(syb7480)* animals displayed a dramatic accumulation of RAD-51 foci in LP cells (Fig. 4A), a phenotype that was previously not reported for the *brc-2(tm1086)*^22^. However, immunostaining for RAD-51 showed that foci accumulate also in the *brc-2(tm1086)*, although at a slightly lower extent (Supp. Fig. 4C). We attribute this discrepancy with previous study^22^ to the use of more sensitive anti-RAD-51 antibodies and improved imaging quality. Quantification of RAD-51 foci in the *ripr-1; brc-2* double mutants show an identical trend of accumulation compared to both *brc-2* and *ripr-1* single mutants (Fig. 4A), with a slight increase in the overall RAD-51 foci number distribution in the *brc-2* and *ripr-1; brc-2* compared to *ripr-1* single mutants. To further assess whether the altered RAD-51 foci profile reflected changes in protein stability, we used the *ripr-1::AID::HA; ieSi38* and *FLAG::AID::brc-2; ieSi38* degron lines crossed into a functional CRISPR-tagged *V5::rad-51*^59^ background. Western blot analyses confirmed that RAD-51 protein stability was unaffected under either RIPR-1 or BRC-2 depletion (Supp. Fig. 4D), further corroborating that the altered foci distribution is due to a genuine loading impairment rather than perturbed protein levels. Together, these results suggest that RIPR-1 and BRC-2 are likely to operate within the same genetic pathway. Moreover, they reveal that contrary to current understanding, RAD-51 can be loaded, at least in LP, independently of BRC-2, unveiling BRC-2-RIPR-1-mediated constraints that are specifically required during early but not later stages of meiotic Prophase I.

### The SC and CO pathway differently modulate RAD-51 loading in ripr-1 and brc-2 mutants

Extensive evidence has shown that under perturbed HR, RAD-51 loading dynamics can be strongly altered, both in terms of foci number and/or distribution across the germ line. For instance, abrogation of CO-mediated DNA repair pathway (e.g. *msh-5* or *cosa-1* mutants)^24,26^ triggers extensive accumulation of RAD-51 in pachytene. Similarly, loss of synapsis results in high numbers of RAD-51 foci that persist in LP^24^. These phenotypes are thought to reflect channeling of the recombination intermediates into inter-sister DNA repair pathways, which depend on the BRC-1/BRD-1 and SMC-5/SMC-6 complexes^60–63^. To test whether elimination of CO pathway affects RAD-51 dynamics in *ripr-1* and *brc-2* mutants, we abrogated *cosa-1* function. To this end, we exploited the *ripr-1::AID::V5; ieSi38* degron strain (Fig. 2), combined with *cosa-1(tm3298)* allele, and analyzed RAD-51 foci upon 24h exposure to auxin, sufficient to reproduce the phenotype of *ripr-1* nulls mutants (Supp. Fig. 4E). Interestingly, depletion of RIPR-1 in the *cosa-1* background led to enhanced RAD-51 accumulation in MP nuclei (Fig. 4B, zone 6). A similar increase was observed in the *cosa-1 brc-2* double mutants (Supp. Fig. 4F, zone 6). In contrast, removal of the SC partially reduced RAD-51 recruitment in LP cells, as observed in *brc-2; syp-3* double mutants (Fig. 4C). These results suggest that removal of pro-CO factors likely channels recombination intermediates into an SC-dependent inter-sister repair pathway(s). Moreover, they reinforce the notion that the constraints imposed by BRC-2-RIPR-1 function on RAD-51 foci formation become less stringent once cells progress through mid-late pachytene stages.

### CO designation is reduced but not abrogated in ripr-1 and brc-2 mutants

Given the highly perturbed RAD-51 dynamics, we next investigated how compromised HR-dependent repair in *ripr-1* and *brc-2* mutants was impacting CO designation. As in most species, CO formation in *C. elegans* is governed by CO interference, a phenomenon by which formation of one CO at a given site prevents the generation of another nearby^64^. Consequently, wild-type meioses typically generate a single CO per homologous chromosome pair, each decorated by several pro-CO factors^25,26,65–68^. Functional tagged lines are available for most of these proteins, and thus we monitored the localization of GFP::MSH-5^61^, GFP::RMH-1^67,69^ and OLLAS::COSA-1^61^ in *ripr-1* and *brc-2* mutants. In whole-mount staining preparations, COSA-1 marks the presumptive CO sites in MP-LP nuclei, forming six discrete foci (corresponding to one CO/homolog pair) that co-localize with MSH-5 and RMH-1^26,67^. At earlier stages (EP), MSH-5/RMH-1 appear as numerous foci, consistent with their recruitment at nascent recombination intermediates.

Loading of pro-CO factors relies upon RAD-51-mediated strand invasion and thus takes place after RAD-51 dissociates from chromatin^26,61,67^. Indeed, analysis of GFP::MSH-5 foci in *ripr-1* mutants (Fig. 5A) showed a stark reduction in zones 2-3 (corresponding to EP-MP), consistent with impaired RAD-51-dependent activity (Fig. 1 and Fig. 3). A similar reduction in GFP::RMH-1 foci was observed in *brc-2* mutants (Fig. 5B). Surprisingly, however, recruitment of GFP::MSH-5, GFP::RMH-1, and OLLAS::COSA-1 at presumptive CO sites in LP nuclei was only partially reduced (∼50%) (Fig. 5A-C) despite the severe impairment of RAD-51 loading at earlier stages, indicating defective but not abrogated CO designation. Co-staining of RAD-51 and COSA-1 revealed that, unlike in control animals, where COSA-1 and RAD-51 rarely overlap^26^, both proteins co-localized extensively in LP nuclei of *ripr-1* and *brc-2* mutants (Supp. Fig. 5A-B). This indicates profoundly altered kinetics in the formation and resolution of the recombination intermediates. These results suggest that pro-CO factors may be recruited at non-canonical, HR-independent repair intermediates, and/or that the residual RAD-51 recruited independently of RIPR-1/BRC-2 in LP cells might be sufficient in promoting MSH-5/RMH-1/COSA-1 loading. To address these hypotheses, we simultaneously abrogated *rad-51* and *ripr-1* function by building the *ripr-1::AID::V5; OLLAS::cosa-1; rad-51(ok2218) ieSi38* strain and quantified COSA-1 foci before and after exposure to auxin. Removal of RAD-51 in RIPR-1-depleted animals reduced COSA-1 foci to comparable levels observed in *rad-51* single mutants (Fig. 5D), indicating that the extensive recruitment of CO-promoting factors observed in absence of RIPR-1 (and likely BRC-2) remains dependent on RAD-51-mediated activity.

**Figure 5.**
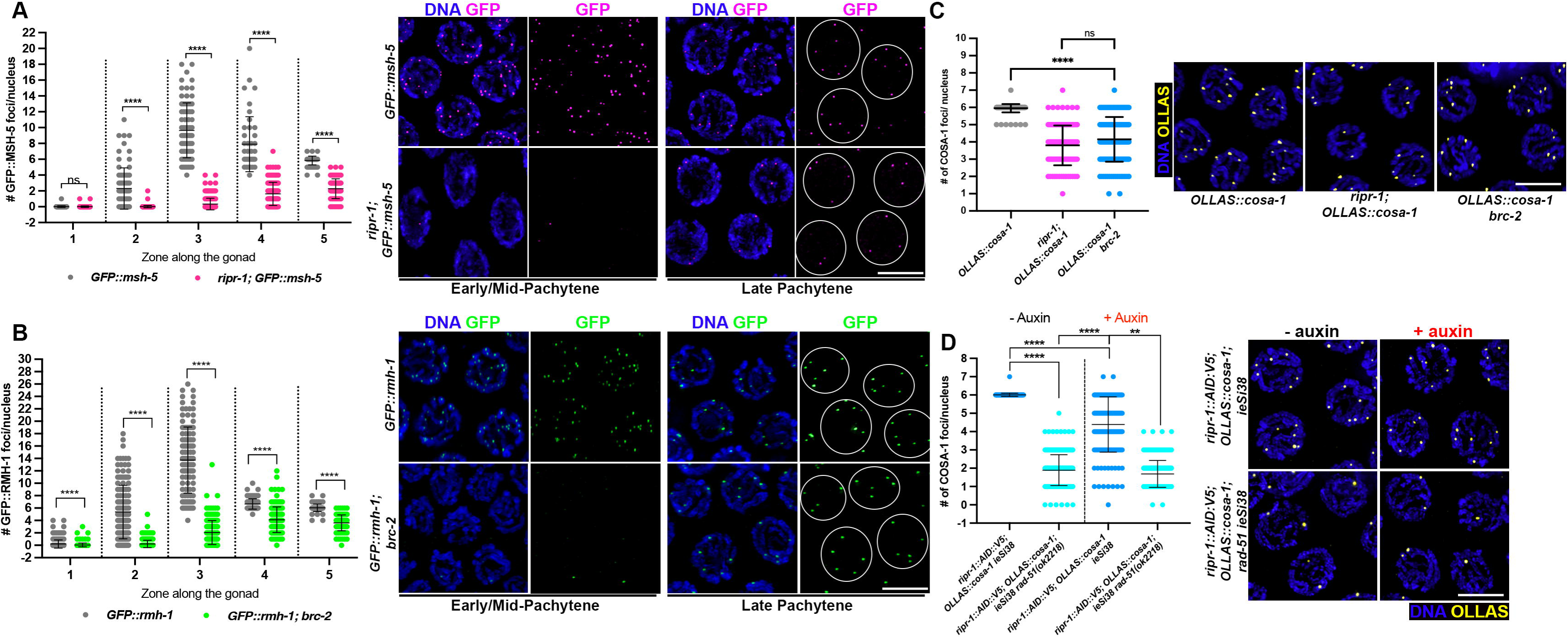
Loss of RIPR-1-BRC-2 reduces but does not abrogate CO designation. **(A)** Left: quantification of GFP::MSH-5 foci number across the germ line in *ripr-1* mutants and control animals. Bars indicate average with S.D., asterisks denote statistical significance as assessed by T test (*\*\*\*\*p<0.0001*). Right: representative images of nuclei from the indicated stages and genetic background immunoassayed for GFP (magenta) and counterstained by DAPI (blue). Circles delineate single nuclei. Scale bar 5 μm. **(B)** Left: quantification of GFP::RMH-1 foci number across the germ line in *ripr-1* mutants and control animals. Bars indicate average with S.D., asterisks denote statistical significance as assessed by T test (*\*\*\*\*p<0.0001*). Right: representative images of nuclei from the indicated stages and genetic background immunoassayed for GFP (green) and counterstained by DAPI (blue). Circles delineate single nuclei. Scale bar 5 μm. **(C)** Left: quantification of OLLAS::COSA-1 foci number in late pachytene cells from *ripr-1* and *brc-2* mutants together with control animals. Bars indicate average with S.D., asterisks denote statistical significance as assessed by T test (*\*\*\*\*p<0.0001*). Right: representative images of nuclei from the indicated stages and genetic background immunoassayed for OLLAS (yellow) and counterstained by DAPI (blue). Scale bar 5 μm. **(D)** Left: quantification of OLLAS::COSA-1 foci number in late pachytene cells from the indicated genetic background and exposure conditions to auxin. Bars indicate average with S.D., asterisks denote statistical significance as assessed by T test (*\*\*p=0.0028*, *\*\*\*\*p<0.0001*). Right: representative images of nuclei from the indicated stages and genetic background immunoassayed for OLLAS (yellow) and counterstained by DAPI (blue). Scale bar 5 μm.

### RIPR-1 and BRC-2 form an obligate complex in vivo and directly interact in vitro

Given their genetic and putative physical interaction, we next wanted to assess whether RIPR-1 and BRC-2 localization is interdependent. To this end, we employed multiple functional CRISPR-tagged alleles for both *ripr-1* (*ripr-1::AID::V5*, *ripr-1::AID::HA* and *ripr-1::GFP*) and *brc-2* (*HA::brc-2*, *FLAG::brc-2* and *FLAG::AID::brc-2*) in different combinations. RIPR-1::AID::V5 displayed a robust nuclear accumulation in both mitotic and meiotic cells, forming puncta/foci structures that did not obviously co-localize with RAD-51 in Pachytene (Fig. 6A), a similar pattern that we have previously shown also for FLAG::BRC-2^28^.

**Figure 6.**
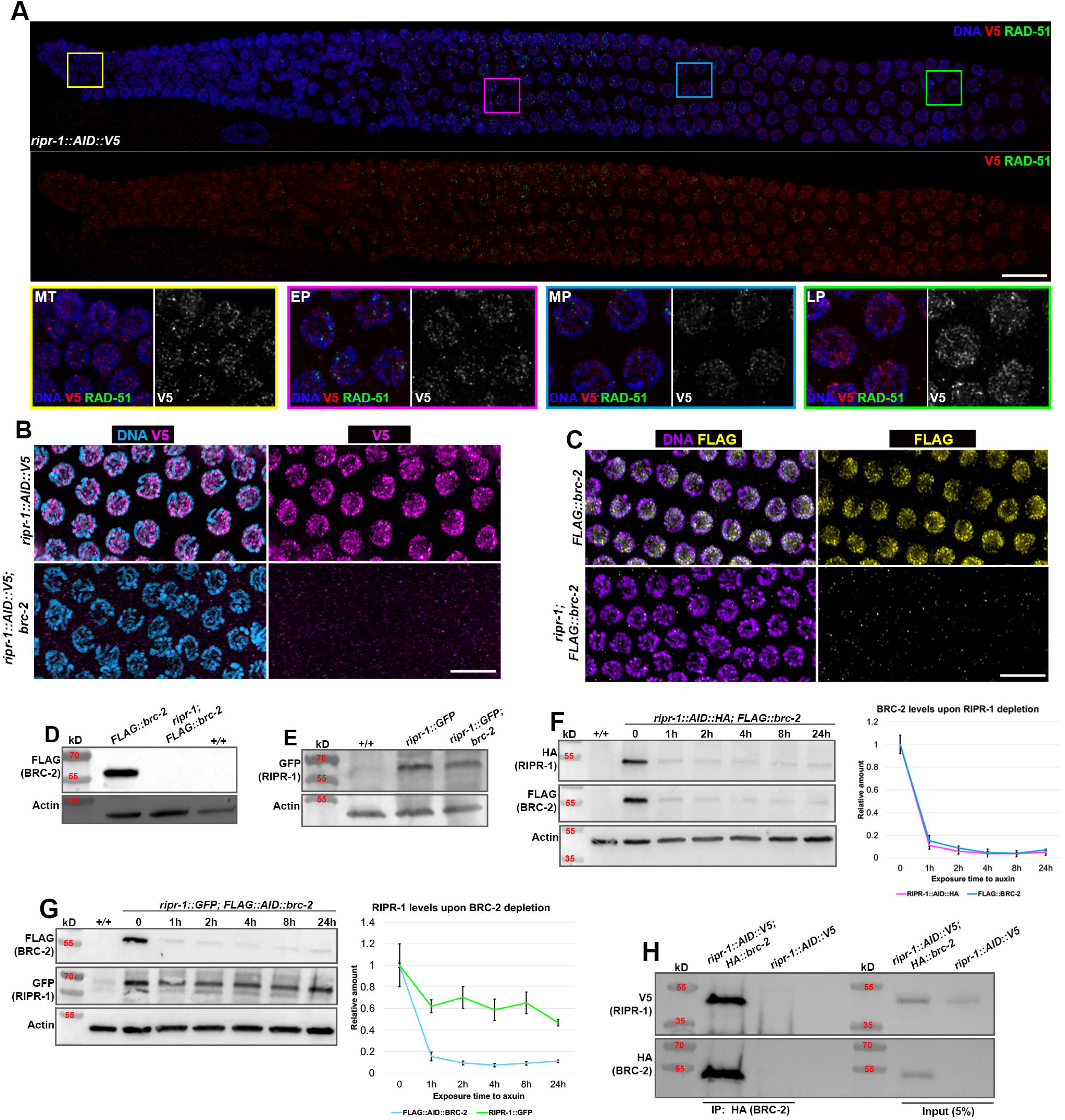
RIPR-1 and BRC-2 form an obligate complex *in vivo*. **(A)** Co-staining of RIPR-1::AID::V5 (red), RAD-51 (green) and DAPI (blue) on whole-mount gonad samples. Colored squares indicate the position along the germ line of the magnified nuclei shown in the insets. Scale bar 5 μm. **(B)** Mid-pachytene nuclei from the indicated genotypes immunoassayed for V5 (RIPR-1, magenta) and DAPI (cyan) in *brc-2* mutants and control animals. Scale bar 5 μm. **(C)** Mid-pachytene nuclei from the indicated genotypes immunoassayed for FLAG (BRC-2, yellow) and DAPI (purple) in *ripr-1* mutants and control animals. Scale bar 5 μm. **(D)** Western Blot on whole cell extracts from the indicated genetic background, probed with anti-FLAG (BRC-2) and anti-Actin antibodies. **(E)** Western Blot on whole cell extracts from the indicated genetic background, probed with anti-GFP (RIPR-1) and anti-Actin antibodies. **(F)** Analysis of RIPR-1::AID::HA and FLAG::BRC-2 total protein levels upon auxin-mediated depletion of RIPR-1 (left) and quantification (right), at the indicated times. Bars indicate S.D. **(G)** Analysis of RIPR-1::GFP and FLAG::AID::BRC-2 total protein levels upon auxin-mediated depletion of BRC-2 (left) and quantification (right), at the indicated times. Bars indicate S.D. **(H)** Western Blot analysis on *in vivo* co-immunoprecipitation assays performed using HA::BRC-2 as bait and relative inputs. BRC-2 robustly and specifically co-immunoprecipitates with RIPR-1.

To assess co-localization, we built the *ripr-1::AID::V5; FLAG::brc-2* strain and performed anti-V5 and FLAG co-staining, which revealed that RIPR-1 and BRC-2 largely overlap in the gonad (Supp. Fig. 6A). Upon ectopic DSB formation by irradiation, both proteins accumulated in discrete chromatin foci in mitotic, but not meiotic cells, and partially co-localized with RAD-51 (Supp. Fig. 6B), reminiscent of BRC-1/BRD-1 behavior^61,62^ and consistent with recruitment to sites of ongoing repair. We also monitored RIPR-1::AID::V5 and HA::BRC-2 expression by Western blot on fractionated protein extracts, which recapitulated their enrichment in the nucleus, as well as displayed low levels of HA::BRC-2 in the cytosol (Supp. Fig. 6C).

Assessment of reciprocal localization revealed a mutual dependent loading, as RIPR-1::AID::V5 signal was lost in *brc-2* mutants (Fig. 6B) and conversely, FLAG::BRC-2 was not detectable in *ripr-1* mutants (Fig. 6C). To distinguish between recruitment or stabilization defect, we analyzed whole cell protein extracts by Western blot. Loss of *ripr-1* resulted in the complete degradation of FLAG::BRC-2 (Fig. 6D), whereas significant levels of RIPR-1::GFP were still detectable in *brc-2* mutants (Fig. 6E). These findings were recapitulated in *ripr-1::AID::V5; brc-2* (Supp. Fig. 6D), *ripr-1::AID::V5; brc-2(tm1086)* (Supp. Fig. 6E) and in *ripr-1::AID::V5; HA::brc-2; ieSi38* exposed to auxin (Supp. Fig. 6F), ruling out potential tag-and/or allele-dependent effects.

To further address the differential stability of RIPR-1 and BRC-2, we gradually depleted each protein using auxin and quantified levels of the non-AID-tagged partner. FLAG::BRC-2 became destabilized as soon as RIPR-1 was reduced (Fig. 6F), whereas ∼50% of the RIPR-1::GFP protein remained detectable upon FLAG::AID::BRC-2 depletion (Fig. 6G).

Together, these results indicate that while each of the proteins is strictly required for the loading of the other in the germ cells, on the other hand BRC-2 stability is strictly dependent on RIPR-1 while RIPR-1 exhibits partial stability in the absence of BRC-2, suggesting an obligate complex with asymmetric dependency.

To confirm their physical interaction, we performed co-immunoprecipitation assays. HA::BRC-2 robustly immunoprecipitated RIPR-1::AID::V5 *in vivo* (Fig. 6H), validating our mass spectrometry results. To independently substantiate their interaction, we undertook two complementary approaches: 1) proximity-labelling using the TurboID biotin ligase fused to a GFP nanobody^70^ targeted to RIPR-1::GFP and followed by mass spectrometry analysis; and 2) we resorted to expression of BRC-2 and RIPR-1 in insect cells to assess direct interaction *in vitro*.

TurboID analysis identified strong enrichment of RIPR-1 compared to control samples, validating our experimental settings (Supp. Table 2 and Supp. Fig. 6G). Crucially, BRC-2 was identified as a strong proximity factor to RIPR-1, consistent with FLAG::BRC-2 pull downs followed by mass spectrometry analysis.

We next assessed whether RIPR-1 and BRC-2 form a stable complex *in vitro* by co-expression and co-purification of both proteins. The identification of both purified 6xHis-RIPR-1 and CBP-BRC-2 was verified by immunoblotting with the corresponding antibody (Supp. Fig. 7A). Notably, co-expression of RIPR-1 markedly improved the solubility of BRC-2 (Supp. Fig. 7B), consistent with the stabilizing effect observed *in vivo* (Fig. 6F and G). Subsequent purification by size-exclusion chromatography revealed formation of a stable RIPR-1–BRC-2 complex of approximately 700 kDa, with apparent 1:1 stoichiometry (Supp. Fig. 7C), suggesting assembly into a higher-order oligomeric complex.

**Figure 7.**
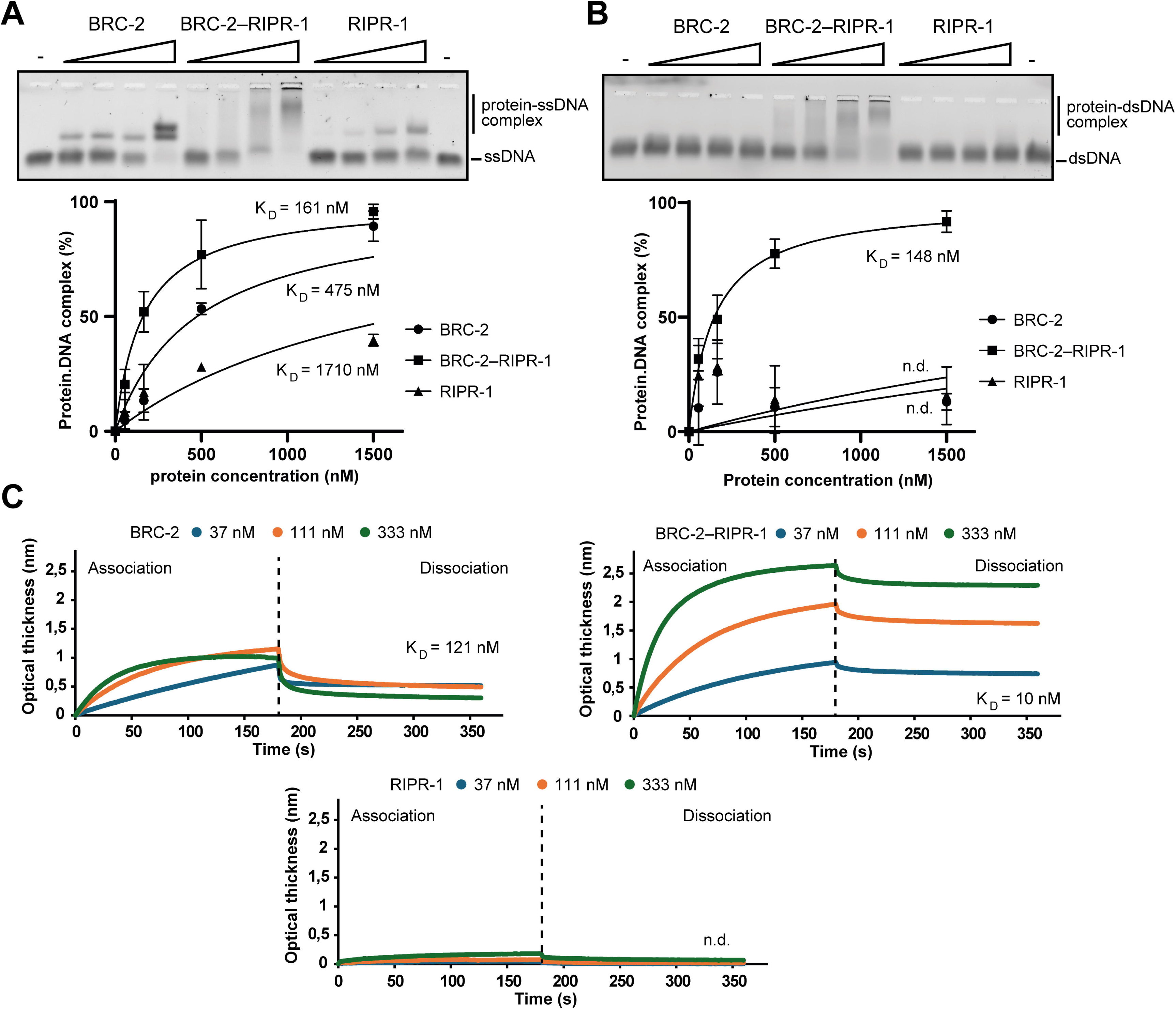
RIPR-1 enhances BRC-2 binding to ssDNA and promotes interaction with dsDNA *in vitro*. Quantification of DNA binding by RIPR-1, BRC-2 and RIPR-1-BRC-2 protein complex to (**A**) ssDNA and **(B)** dsDNA substrates, assessed by EMSA with corresponding representative agarose gel. Increasing concentrations of purified proteins were incubated with fluorescently labelled DNA substrates prior to electrophoretic separation. **(C)** DNA-binding kinetics of BRC-2, RIPR-1-BRC-2 and RIPR-1 by BLI assay using immobilized biotinylated ssDNA substrate. Representative sensor grams show real-time protein association with surface-bound ssDNA (association phase) followed by transfer of sensors into protein-free buffer to monitor dissociation kinetics (dissociation phase).

Taken together, our data show that RIPR-1 and BRC-2 undergo a mutually dependent loading in the germ line, they establish an obligate complex *in vivo* and directly associate *in vitro*.

### RIPR-1 enhances the DNA binding affinity of BRC-2

The ability of purifying both individual proteins and the RIPR-1–BRC-2 complex (Supp. Fig. 7A) enabled us to investigate the impact of RIPR-1 on DNA binding. Given that RIPR-1 contains a predicted canonical OB-fold domain (Fig. 1), we first assessed its intrinsic binding to ssDNA using electrophoretic mobility shift assays (EMSA). Gel filtration analysis revealed that RIPR-1 elutes as two distinct species (∼ 670 and ∼ 120 kDa, Supp. Fig. 7C), which were tested separately for binding to a dT79 ssDNA substrate. Notably, only the higher-molecular-weight (HMP) complex exhibited ssDNA binding (Supp. Fig. 7D), albeit weaker than that of BRC-2 (Fig. 7A), whereas the lower-molecular-weight form (LMP) showed no detectable interaction under tested conditions (Supp. Fig. 7D).

Incorporation of RIPR-1 into the complex with BRC-2 enhanced DNA binding compared with BRC-2 alone (Fig. 7A), suggesting a functional contribution of RIPR-1 to nucleoprotein complex stability. Strikingly, while neither RIPR-1 nor BRC-2 alone displayed detectable binding to dsDNA, the RIPR-1/BRC-2 complex showed robust dsDNA-binding (Kd ∼148 nM), comparable to its interaction with ssDNA (Fig. 7B). This observation suggests that the complex formation alters the DNA-binding properties of BRC-2, potentially broadening its substrate specificity or stabilizing transient interactions with DNA structures containing both ss- and dsDNA regions.

To quantitatively assess these effects, we employed biolayer interferometry (BLI) using an immobilized biotinylated dT80 ssDNA substrate. Consistent with the EMSA results, RIPR-1 alone displayed low affinity for ssDNA (Fig. 7C, bottom), whereas BRC-2 bound with high affinity (Kd ∼121 nM) (Fig. 7C, top left). Strikingly, the RIPR-1–BRC-2 complex exhibited an approximately 12-fold increase in affinity (Kd ∼10 nM, Fig. 7C, top right). Kinetic analysis revealed that this enhancement is primarily driven by a reduced dissociation rate, indicating that RIPR-1 stabilizes BRC-2 binding to ssDNA.

We next examined binding to dsDNA by BLI. In agreement with EMSA, neither BRC-2 nor RIPR-1 individually showed detectable interaction (Supp. Fig. 7E, top and bottom respectively) whereas RIPR-1–BRC-2 complex bound dsDNA with enhanced affinity (Kd ∼123 nM, Supp. Fig. 7E, middle).

Collectively, these data demonstrate that RIPR-1 potentiates the DNA-binding activity of BRC-2 and broadens its substrate specificity, likely by stabilizing its interaction with DNA. This provides a mechanistic basis for their cooperative function during homologous recombination.

### Loading of the RIPR-1–BRC-2 complex does not require interaction with RAD-51

Previous studies *in vitro* demonstrated that BRC-2 directly interacts with RAD-51 through its conserved BRC motif, and that this association is essential to promote D-loop formation^22,71^. Specifically, RAD-51 binds two conserved BRC-2 residues, Phe35 (F35) and Ile43 (I43)^72^, and mutations of either residue abrogate pro-recombinogenic activity without preventing BRC-2 binding to ssDNA^22,71,72^. However, the functional relevance of F35/I43 to BRC-2 activity *in vivo* has not been assessed.

Co-immunoprecipitation experiments confirmed that BRC-2 and RAD-51 form a complex *in vivo* under our experimental conditions (Fig. 8A), consistent with previous report^22^. To investigate the contribution of RAD-51 binding to BRC-2 function, we generated the *FLAG::brc-2^F35A^* mutant line by CRISPR and analyzed viability, protein stability, and RAD-51 localization. *FLAG::brc-2^F35A^* mutant worms displayed complete embryonic lethality (Fig. 8B), and RAD-51 immunolabelling showed dramatic accumulation of recombination intermediates in MP-LP cells (Fig. 8C-D), recapitulating *brc-2* null mutants. Cytological detection of FLAG::AID::BRC-2^F35A^ revealed normal nuclear localization in the germ line (Fig. 8E), and Western blot analysis on whole-cell extracts showed only a slight, although not significant, reduction of BRC-2^F35A^ levels coupled with reduced protein mobility (Fig. 8F). These results indicate that RAD-51 binding is dispensable for BRC-2 recruitment *in vivo* and overall stability, in line with the results obtained *in vitro*^22,71^. Furthermore, we monitored whether disruption of the RAD-51-BRC-2 interaction affected RIPR-1 recruitment. Localization of *ripr-1::GFP* in *FLAG::AID::brc-2^F35A^*worms appeared unaltered, with no evident changes in protein loading or expression (Fig. 8G). Collectively, these results indicate that binding to RAD-51 is not required for RIPR-1-BRC-2 complex recruitment. Importantly, they also reveal that recruitment of RAD-51 to meiotic recombination intermediates can occur independently of its binding to BRC-2 during mid/late-pachytene stage, unveiling novel and unanticipated mechanisms that drive execution of HR.

**Figure 8.**
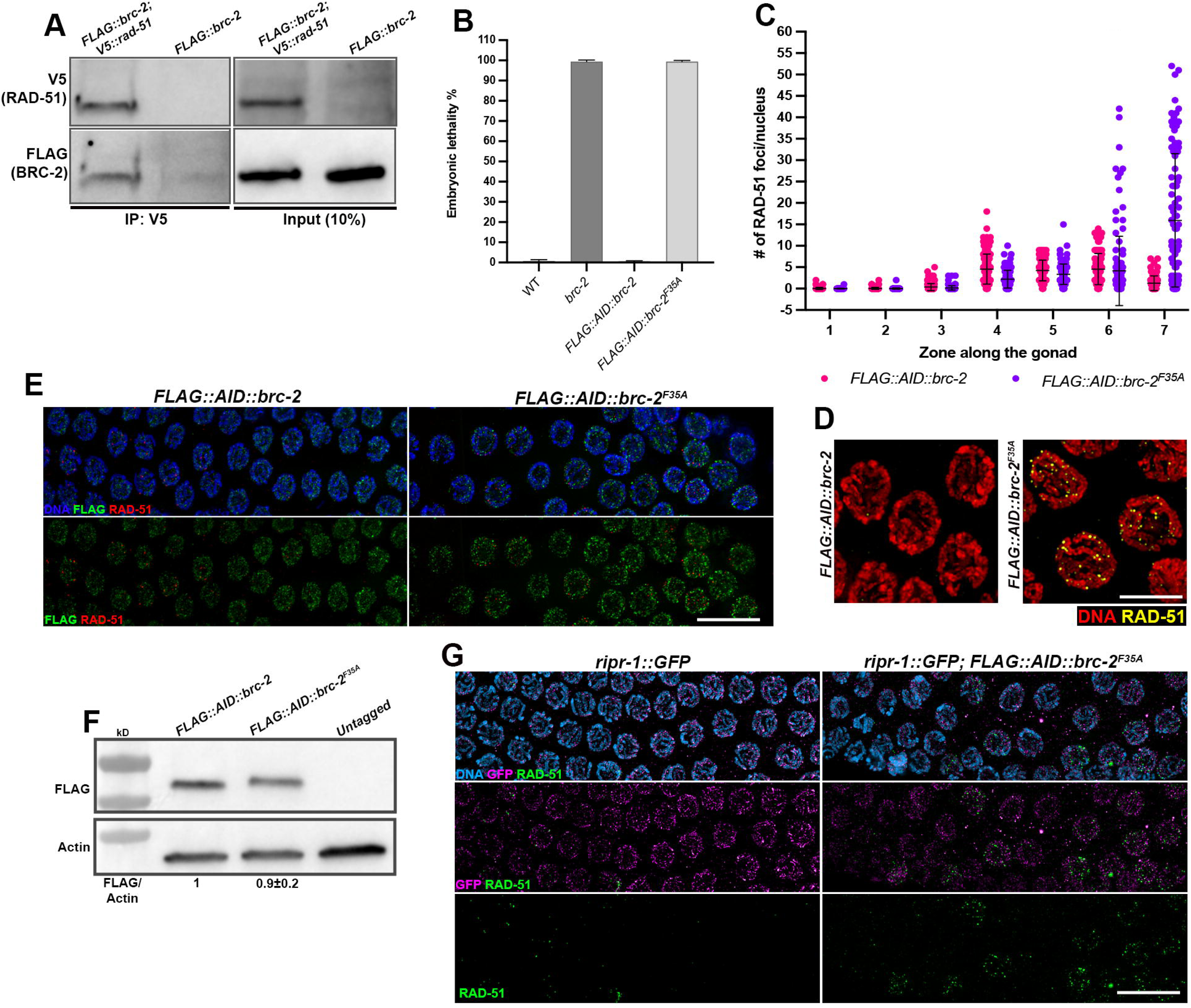
Binding to RAD-51 is not essential to recruit RIPR-1-BRC-2 to chromatin *in vivo*. (A) *In vivo* co-immunoprecipitation of FLAG::BRC-2 and V5::RAD-51. **(B)** Assessment of embryonic lethality in the indicated genotypes. Bars depict average with S.D. **(C)** Quantification of RAD-51 foci across the germ line of *FLAG::AID::brc-2* control animals and *FLAG::AID::brc-2^F35A^*mutants. **(D)** Representative images of late pachytene nuclei from the indicated strains stained for RAD-51 (yellow) and counterstained by DAPI (red). Scale bar 5 μm. **(E)** Late pachytene snapshots from the indicated genotypes immunoassayed for RAD-51 (red) and FLAG (BRC-2, green) and counterstained by DAPI (blues) showing comparable loading of BRC-2F35A as BRC-2^WT^. Scale bar 5 μm. **(F)** Western blot on whole-cell extracts showing expression of FLA::AID::BRC-2^F35A^ protein. Actin was used as loading control and FLAG/Actin ratio is shown on the bottom. **(G)** Late pachytene snapshots from the indicated genotypes immunoassayed for RAD-51 (green) and GFP (RIPR-1, magenta) showing normal localization of RIPR-1 in the *FLAG::AID::brc-2^F35A^*mutant animals. Scale bar 5 μm.

## DISCUSSION

We identified RIPR-1 as a previously uncharacterized regulator of meiotic homologous recombination and show that it forms an obligate complex with BRC-2/BRCA2 in *C. elegans*. This complex is required for efficient RAD-51 loading, timely processing of recombination intermediates, crossover designation, and chromosome integrity. Identification of RIPR-1 advances our understanding of the molecular network supporting BRC-2 activity, which, until now, remained entirely uncharacterized beyond the direct BRC-2–RAD-51 interaction^22^.

### RIPR-1 as a structural regulator of BRC-2 activity

In mammals, BRCA2 cooperates with several accessory factors, including PALB2/FANCN, which loads BRCA2 to sites of damage^73–75^, and meiosis-specific proteins MEILB2 and BRME1^19,20^. By contrast, clear homologues of these BRCA2 co-factors are not readily identifiable outside vertebrates, significantly hindering our understanding of how BRCA2-like proteins are regulated in simpler model organisms. Identification of RIPR-1 partially fills this gap in *C. elegans*, providing an entry point to understanding BRC-2 cofactors networks in a genetically tractable metazoan system. RIPR-1 and BRC-2 establish an obligate complex both *in vitro* and *in vivo*, with an asymmetric dependency: BRC-2 stability entirely depends on presence of RIPR-1, while the latter retains partial stability in the absence of BRC-2. This architecture resembles the MEILB2-BRME1 relationship, in which BRME1 stabilizes MEILB2 within the ternary BRCA2-MEILB2-BRME1 complex^19,20^. Consistent with a structural role, co-expression of RIPR-1 markedly improved BRC-2 solubility and the two proteins co-purify as a stable ∼700 kDa complex with apparent 1:1 stoichiometry, suggesting assembly into a higher-order oligomeric state.

We confirm that RIPR-1-BRC-2 acts downstream DNA end resection but upstream of productive RAD-51 filament formation. In *ripr-1* mutants, MRN/X localization and robust RPA-1 accumulation indicate that ssDNA is generated and stabilized, yet RAD-51 loading is severely impaired during early meiotic prophase. These data place RIPR-1–BRC-2 complex within the conserved class of recombination mediators that promote the transition from RPA-coated ssDNA to RAD-51 nucleoprotein filaments. Such mediator activity is central to HR because RPA binds ssDNA with high affinity and must be displaced or remodeled to allow assembly of RAD-51 filament. RIPR-1 harbors a predicted N-terminal OB-fold domain, a versatile structural motif found in many genome guardians, such as BRC-2 itself, RPA-1, and RMH-1^29,30,76,77^, which confers conformational flexibility essential for protein sliding along DNA, protection of ssDNA from nucleolytic degradation, and dynamic regulation of DNA binding within multiprotein complexes. This might explain why RIPR-1 potentiates the DNA-binding activity of BRC-2 enhancing ssDNA affinity of approximately 12-fold, primarily by reducing the dissociation rate and enabling robust dsDNA binding that is absent in either protein alone. This differs from human BRCA2, that achieves both ssDNA and dsDNA engagement through its intrinsic C-terminal OB-fold-containing DNA-binding domain (DBD)^78^. *C. elegans* BRC-2 lacks an equivalent extended DBD, and the gain of dsDNA binding upon complex formation suggests that RIPR-1 compensates for this deficiency. Notably, in the human system, the BRC repeats of BRCA2 play a dual role: they target RAD51 to ssDNA and concurrently prevent its nonproductive nucleation onto dsDNA^79,80^. This ssDNA/dsDNA selectivity is essential to ensure that RAD51 is directed specifically to resected DSB ends rather than engaging intact duplex DNA.

Our work here shows that *C. elegans* BRC-2 alone cannot engage dsDNA, and this capacity is only acquired upon RIPR-1 complex formation. We speculate that RIPR-1 may therefore contribute not only to RAD-51 loading efficiency but also to regulating the substrate preference of the complex at resection junctions that contain both ssDNA tails and adjacent dsDNA. This architecture is reminiscent of the DSS1-BRCA2 partnership where DSS1 promotes RPA-RAD51 exchange by engaging RPA and modulating BRCA2 function^81^, and additionally restrains excessive BRCA2 engagement with dsDNA to favor productive RAD51 loading onto ssDNA^86^. RIPR-1 is unlikely to be a functional DSS1 equivalent, but represents a distinct, nematode-specific solution that stabilizes BRC-2 and broadens its DNA-binding competence, supporting the broader principle that BRCA2-family proteins rely on accessory factors to tune their stability, substrate preference, and recombination mediator activity.

A central and unexpected finding of our work is that RAD-51 loading is not strictly dependent on BRC-2 function throughout meiotic Prophase I. Consistent with previous data, we show that indeed loss of BRC-2, either directly or by abrogating RIPR-1 function, severely impairs RAD-51 foci formation during early-to-mid pachytene, as similarly observed in other organisms^82,83^. Strikingly however, we found that this does not hold true in mid- and late pachytene cells, in which abrogation of RIPR-1–BRC-2 complex function triggers a massive formation of chromatin-associated RAD-51 foci, whose recruitment requires EXO-1-DNA-2-dependent long-range resection^54^. Critically: (i) these recombination intermediates originate from SPO-11-dependent DSBs; (ii) they are independent of apoptotic-induced DNA damage; (iii) they can sustain substantial levels of CO designation through RAD-51-mediated activity; and (iv) they arise also in the *brc-2^F35A^* point mutant that disrupts RAD-51 binding to the BRC motif, confirming that alternative loading is mechanistically independent of the canonical BRC-2-RAD-51 interface. These results collectively indicate that BRC-2 activity is not required to allow RAD-51 loading *per se* but rather constrains its recruitment at resected DSBs in a cell-stage-dependent fashion.

### How is a BRC-2 independent RAD-51 loading achieved in C. elegans?

We envision three non-mutually exclusive scenarios for this alternative loading. The first posits that an as-yet-unidentified mediator may substitute for BRC-2–RIPR-1 activity specifically in late pachytene cells.

The second invokes the intrinsic dynamics of RPA combined with changes in chromatin context. Recent single-molecule studies have established that RPA’s OB-fold domains undergo rapid “breathing”, a transient microscopic dissociation from ssDNA that intermittently exposes short stretches of ssDNA between neighboring RPA molecules^84–86^. Recombination mediators exploit this breathing to gain access to ssDNA and initiate RAD-51 filament nucleation^85,86^. Crucially, free RPA inhibits RAD-51 nucleation but has a comparatively lesser impact on filament elongation^84^, suggesting that once a nucleation event occurs, RAD-51 filament growth can proceed even in the presence of competing RPA.

In the absence of an active BRCA2-type mediator, extended ssDNA substrates generated by long-range resection could, in principle, provide sufficient transient access *via* RPA OB-fold breathing, for RAD-51 nucleation to occur spontaneously. This passive model is consistent with our observation that EXO-1/DNA-2-dependent generation of long ssDNA tracts is an obligate prerequisite for late RAD-51 accumulation in *ripr-1* and *brc-2* mutants: longer ssDNA substrates increase the statistical probability of transiently exposed nucleation sites between RPA molecules.

The third scenario concerns the loss of BRC-2-mediated suppression of RAD-51 loading onto dsDNA. In the human system, BRC repeats not only promote RAD51 assembly on ssDNA but also actively prevent its nonproductive nucleation on dsDNA, thereby directing the recombinase specifically to resected ends^79,80^. In the *C. elegans* context, BRC-2 is likely to exert an analogous restraint *via* its BRC motif. When BRC-2 is absent, this restraint is relieved, potentially allowing RAD-51 to nucleate on the dsDNA flanking resection tracts. This model is consistent with our *in vitro* observation that the RIPR-1-BRC-2 complex uniquely acquires dsDNA-binding capacity, a property we propose may serve to occupy dsDNA at resection junctions and spatially restrict RAD-51 from engaging the adjacent duplex. In its absence, unregulated RAD-51-dsDNA association may generate ectopic, non-productive filaments that appear as RAD-51 foci cytologically but fail to support faithful inter-homolog repair. This would explain the paradox at the heart of our findings: robust RAD-51 recruitment in late pachytene, yet severely compromised Diakinetic chromosomes. We note that this scenario is not mutually exclusive with the OB-fold breathing model; both could operate simultaneously in LP cells deprived of BRC-2–RIPR-1, with the relative contribution of ssDNA-based passive loading versus ectopic dsDNA loading potentially influenced by the extent of resection and chromatin accessibility.

The stage-specificity of this alternative loading is likely further shaped by changes in chromosome architecture. Germ cells in the *C. elegans* gonad switch between different modes of DSB repair as they progress through meiotic prophase, a transition that is in part likely dependent on changes of chromosome structure^56^. Chromatin spatial configuration in meiotic cells appears to be globally more constrained into a reduced nuclear area in transition zone and early pachytene compared to mid- and late pachytene stages, where the homologs appear rather individualized and with a linear morphology. These changes in chromosome architecture are actively regulated and are key to proper execution of different meiotic tasks, including a strict control of RAD-51 foci formation and repair^87^. It is therefore plausible that the more “open” chromatin architecture in late pachytene, combined with the extended ssDNA generated by long-range resection, lowers the threshold for “passive” recruitment of RAD-51 along resected DSBs in late pachytene cells, rendering BRC-2–RIPR-1-mediated active loading dispensable while simultaneously removing the safeguard against nonproductive filament formation.

Our analysis of CO-designation markers shows that this late BRC-2-independent RAD-51 recruitment is required to maintain residual loading of MSH-5-RMH-1-COSA-1 through a RAD-51-dependent mechanism, as directly demonstrated by the epistatic reduction of COSA-1 foci upon simultaneous loss of RAD-51 and RIPR-1. The concomitant accumulation of RAD-51 and COSA-1 in late pachytene, contrasting their non-overlapping distribution in wild-type animals, reflects profoundly altered kinetics of recombination intermediate formation and resolution. However, we cannot formally exclude that a fraction of these pro-CO factors are recruited at non-canonical recombination events, given that MSH-5, RMH-1, and COSA-1 have been reported to accumulate along chromosomes under dysfunctional HR, synapsis and loss of cohesins^69,88,89^. Nonetheless, the RAD-51-dependence of COSA-1 loading argues that at least a fraction reflects genuine, albeit delayed and aberrant, CO designation events driven by the alternative recombinase loading pathway.

## Conclusions

Viewed in broader evolutionary perspective, this work highlights how the modular architecture of BRCA2-dependent RAD51 loading can diversify while preserving its core function. Human BRCA2 encodes three OB folds and a helical domain within its C-terminal DBD that directly engages DNA substrates^84^, so that ssDNA/dsDNA engagement and the dual regulation of RAD51 recruitment and exclusion from dsDNA are all intrinsic to a single polypeptide.

*C. elegans* BRC-2 is comparatively reduced in this domain and requires RIPR-1 to achieve full DNA-binding competence and recombination mediator activity. The biological system thus resolves the same problem - efficient, substrate-specific recombinase loading - through a two-subunit complex rather than a single extended protein. Analogously, while the DSS1-BRCA2 interaction fine-tunes substrate engagement in human cells^81,90^ the RIPR-1–BRC-2 interaction achieves a related but non-equivalent outcome through a distinct structural partnership. These observations support a broader model in which BRCA2-family proteins can evolve compositionally different solutions to the recombination mediator problem, preserving the essential activity of promoting recombinase assembly on RPA-coated ssDNA at resected DSBs.

Together, our work identifies RIPR-1 as a critical BRC-2 factor that stabilizes this recombinase loader complex and expands its DNA-binding repertoire substrates. It reveals a previously unanticipated layer of spatiotemporal regulation in the meiotic recombination program where BRC-2–RIPR-1-mediated constraints on RAD-51 loading are specifically operative during early meiotic prophase and are progressively relaxed as cells approach late pachytene, at which point RAD-51 can access resected DNA through a BRC-2-independent pathway, possibly facilitated by the intrinsic breathing dynamics of RPA OB-folds on extensively resected ssDNA substrates. This unexpected plasticity in recombinase assembly demonstrates that the pathway from DSB resection to strand-exchange competent RAD-51 filaments is more nuanced than previously appreciated and establishes the *C. elegans* germ line as a valuable system to dissect the spatiotemporal logic of BRCA2-dependent DNA repair.

## METHODS

### Genetics

All strains were maintained on NG medium plates seeded with OP50 *E. coli* as food source. All experiments were conducted at 20°C and a full list of the strains used in this study is provided in the Supplementary Table 3. All genetic edits were made by Suny Biotech (https://www.sunybiotech.com/) by CRISPR/Cas9 to generate knock-in, knock-out or point mutations, unless previously generated and already available for study. Analyses of viability were performed in biological duplicates by selecting individual L4 animals of the relevant genotypes, that were transferred onto fresh plates every 24h until unfertilized eggs were laid. The embryonic lethality was calculated as the total number of dead eggs/total number of laid eggs.

### Immunofluorescence studies

Synchronized animals from the relevant mutant strains and controls at the indicated age and exposure conditions to auxin were dissected as previously described without modifications^43^. For detection of GFP::MSH-5, worms were picked at L4 stage and incubated at 20°C for 20-24 hours. They were then dissected in 1×PBST on poly-l-lysine-coated SuperFrost slides, immediately freeze-cracked in liquid nitrogen, incubated in methanol at −20°C for 20 minutes, and then incubated in fixation buffer (100 mM phosphate buffer containing 4% PFA) at room temperature for 15 minutes. After three 5-minutes washes in 1×PBST, the samples were incubated in 4’,6-diamidino-2-phenylindole (DAPI) (2 μg/ml) for 1 minute. Finally, the slides were washed once in 1×PBST for 20 minutes and mounted with Vectashield. A complete list of antibodies used in this study is provided in Supplementary Table 4.

Analyses of DAPI bodies in Diakinesis nuclei were performed only in the most proximal nuclei (−1 and −2 oocytes), and the number of nuclei assessed for each experiment is reported in Supplementary Table 5.

Quantification of RAD-51 foci number was performed as previously described. Briefly, gonads were divided into seven equal regions from the mitotic tip to Diplotene entry. The number of foci was scored for each nucleus and the distribution was plotted.

The same procedure was followed to quantify synapsis, but the gonads were divided into six equal regions starting with the TZ through entry into Diplotene.

The same zoning was applied to quantification of GFP::MSH-5 and GFP::RMH-1 foci, whereas the number of COSA-1 foci was quantified only in the last seven cell rows starting from the first row of nuclei before the gonadal bend going backwards.

The number of nuclei analyzed for RAD-51, GFP::MSH-5, GFP::RMH-1 and OLLAS::COSA-1 foci quantification is reported in the Supplementary Table 5.

Collection of fixed images was performed using a fully motorized widefield upright microscope Zeiss AxioImager Z2, equipped with a monochromatic camera Hamamatsu ORCA Fusion, sCMOS sensor, 2304 x 2304 pixels, 6.5 x 6.5 μm size. Z-stacks were set at 0.25 μm thickness and images were deconvolved using ZEN Blue Software using the “constrained iterative” algorithm set at maximum strength. Whole projections of deconvolved images were generated with Fiji (ImageJ) and processed in Photoshop, where some false colouring was applied.

Imaging of GFP::MSH-5 was performed with a DeltaVision Ultra microscope (100× oil immersion objective lens; N.A. 1.4) equipped with a cMOS (complementary metal oxide semiconductor) camera. Stacks of 0.2 µm were taken and deconvolved with SoftWoRx software using the conservative mode coupled with 15 iterations.

### Biochemistry

To generate whole-cell protein extracts, 100 animals from each genotype and exposure conditions to auxin were picked in 1x Tris EDTA (pH 7) containing 1x protease inhibitor (Roche). Samples were snap frozen in liquid nitrogen and quickly thawed before adding Laemmli Buffer to 1x final concentration. Samples were boiled for 10 minutes, briefly spun down and loaded on 4-20% precast acrylamide gels (Biorad). Gels were run in 1x Tris-Glycine buffer containing 0.1% SDS and transferred onto a nitrocellulose membrane for 90 minutes at 100V in 1x Tris-Glycine buffer containing 20% methanol at room temperature. Membranes were rinsed in 1x TBST (1x TBS with 0.1% Triton) and blocked in 1x TBST containing 5% non-fat milk. Primary and secondary antibodies were added in the same buffer and probed overnight at 4°C and 2h at room temperature respectively.

For immunoprecipitation assays, large cultures of synchronized worms were employed to produce fractionated nuclear extracts as in^91^ without modifications. 1-2 mg of nuclear extract (nuclear soluble and chromatin-bound pooled together) were incubated with 40 μl of agarose-magnetic V5, HA or agarose GFP traps (Proteintech) for 4 hours at 4°C in buffer D (20 mM HEPES pH 7.6, 200 mM KCl, 1 mM EDTA pH 8, 20% glycerol, 0.2% Triton and 1x cOmplete protease inhibitor). Beads were recovered with a magnetic stand or by centrifugation (7500 rpm for 5 minutes) and washed extensively with buffer D. Immunocomplexes were eluted from beads by adding 40 μl of 2x Laemmli Buffer and samples were boiled for 10 minutes. Beads were separated by centrifugation and the entire supernatant was loaded on a 4-20% acrylamide gel and processed as above.

The blots were developed by Clarity Max ECL (BioRad) and images were acquired using the Azure 400 Western Blot Imager.

Samples for Western blot shown in Supp. Fig. 7 were prepared in SDS buffer (0.16 M Tris HCl pH 6.8, 4% SDS, 20% glycerol, 0.01% bromophenol blue and 100 mM DTT), separated on 12% SDS-PAGE at 150 V for 60 min, and proteins transferred to nitrocellulose membrane using the semi-dry Trans-blot turbo Transfer system (1704150; Biorad). The membrane was blocked with 5% milk in PBS for 60 min at room temperature, followed by overnight incubation at 4 °C with primary antibodies: rabbit anti CBP (05-932 Sigma-Aldrich, 1:2000 in TBST with 1% BSA), rabbit HRP-conjugated anti GAPDH (3683S, Cell Signalling, 1:1000 in 5% milk), mouse anti polyHis (H1029, Sigma-Aldrich, 1:2000 in 5% milk), rabbit anti vinculin (ab129002, Abcam, 1:1000 in 5% milk). After washing, the membrane was incubated with Anti-rabbit IgG, HRP-linked Antibody (A6154, Sigma-Aldrich, 1:5000 in 5% milk) or Anti-Mouse IgG–Peroxidase antibody (A0168, Sigma-Aldrich, 1:5000 in 5% milk) for 60 min at room temperature. The blots were developed by the Immobilon Western Chemiluminescent HRP Substrate (WBKLS0500; MERCK Millipore), and images were acquired using the Luminescent Image Analyser (ImageQuant™ LAS 4,000; Fujifilm).

### Mass Spectrometry on FLAG::BRC-2 pulldowns

FLAG pull downs were performed by using M2 FLAG magnetic beads (Sigma, #M8823) on fractionated nuclear protein extracts produced from *FLAG::brc-2* and WT untagged negative controls. Immunocomplexes were eluted from beads by adding 40 μl of 2x Laemmli buffer, followed by boiling for 10 minutes. Samples were spun down, loaded onto a pre-cast 4-20% acrylamide gradient gel (BioRad) and run for approximately 15 minutes at 150 V in 1x Tris-Glycine buffer containing 0.1% SDS at room temperature. Gel was stained with Coomassie Brilliant Blue and destained in bi-distilled water for 1 hour.

1D gel lanes were excised manually and after destaining and washing procedures each band was subjected to protein reduction (10mM DTT in 25mM NH_4_HCO_3_, 45 min, 56°C, 750 rpm) and alkylation (55mM IAA in 25mM NH_4_HCO_3_; 30 min, laboratory temperature, 750 rpm) step. After further washing by 50% ACN/NH_4_HCO_3_ and pure ACN, the gel pieces were incubated with 125 ng trypsin (sequencing grade; Promega) in 50mM NH_4_HCO_3_. The digestion was performed overnight at 37 °C on a Thermomixer (750 rpm; Eppendorf). Tryptic peptides were extracted into LC-MS vials by 2.5% formic acid (FA) in 50% ACN with addition of polyethylene glycol (final concentration 0.001%)^92^ and concentrated in a SpeedVac concentrator (Thermo Fisher Scientific).

LC-MS/MS analyses were done using RSLCnano system connected to Orbitrap Exploris 480 spectrometer (Thermo Fisher Scientific) with EASY Spray ion source (Thermo Fisher Scientific) installed. Prior to LC separation, tryptic digests were online concentrated and desalted using trapping column (300 μm × 5 mm, μPrecolumn, 5μm particles, Acclaim PepMap100 C18, Thermo Fisher Scientific). After washing of trapping column with 0.1% FA, the peptides were eluted (flow 300 nl/min) from the trapping column onto Acclaim PepMap RSLC C18 column (2 µm particles, 75 μm × 500 mm; Thermo Fisher Scientific) by 108 min long gradient (mobile phase A: 0.1% FA in water; mobile phase B: 0.1% FA in 80% acetonitrile).

Data were acquired in a data-independent acquisition mode (DIA). The survey scan covered m/z range of 350-1400 at resolution of 60,000 (at m/z 200) and maximum injection time of 55 ms. HCD MS/MS (27% relative fragmentation energy) were acquired in the range of m/z 200-2000 at 30,000 resolution (maximum injection time 55 ms). Overlapping windows scheme in m/z range from 400 to 800 were used as isolation window placements.

DIA data were processed in DIA-NN^2^ (version 1.8) against modified cRAP database (based on http://www.thegpm.org/crap/, 112 sequences in total) and UniProtKB protein database for *Caenorhabditis elegans* (number of protein sequences: 26,601). No optional, carbamidomethylation as fixed modification and trypsin/P enzyme with 1 allowed missed cleavages and peptide length 7-30 were set during the library preparation. False discovery rate (FDR) control was set to 1% FDR. MS1 and MS2 accuracies as well as scan window parameters were set based on the initial test searches (median value from all samples ascertained parameter values). MBR was switched on.

Intensities of reported proteins were further evaluated using software container environment (https://github.com/OmicsWorkflows; version 4.1.3a). Processing workflow is available upon request. Briefly, it covered: a) removal of low-quality precursors and contaminant protein groups, b) protein group intensities log2 transformation, LOESS normalization, c) ratio calculation of protein group intensities.

### Nuclei collection and pull down for Turbo-ID mass spectrometry analysis

Synchronized adult worms (24h post-L4) were collected in a buffer containing 10mM HEPES-KOH pH 7.6, 1mM EDTA (pH 8), 10mM KCl, 1.5mM MgCl_2_, 0.25mM Sucrose, 1mM PMSF and 1mM DTT, snap-frozen in liquid nitrogen and stored at −80°C until further use. After defrosting, the worms were dounced and the solution obtained was filtered through a 100µm and then a 40µm cell strainer. The final flow-through was centrifuged at 300g for 2 minutes at 4°C. The supernatant was then centrifuged at 2500g for 10 minutes at 4°C. The pellet was resuspended in nuclear lysis (NL) buffer supplemented with protease inhibitor and DTT (obtained from Qproteome nuclear protein kit, catalog # 37582) and incubated on ice for 15 minutes after which NP buffer (obtained from the same kit) was added. This was centrifuged at 10000g for 5 minutes at 4°C. The pellet was resuspended in a buffer containing 6M Guanidinium, 50mM Tris (pH 8), 100mM NaCl, 5mM DTT and Protease Inhibitor cocktail (1 tablet/10mL, (cOmplete™, Mini, EDTA-free Protease Inhibitor Cocktail, catalog #11836170001)) and centrifuged at 13000g for 20 minutes at room temperature. The supernatant was added to 20µl of Streptavidin magnetic beads (Pierce™ Streptavidin Magnetic Beads, catalog #88816) that were equilibrated using Solution 1 (50mM Tris pH 7.5 and 100mM NaCl) and incubated at room temperature on a rotating wheel for one hour. The beads were washed four times with Solution 2 (6M Guanidinium, 50mM Tris pH 8, 100mM NaCl), four times with Solution 1, and finally washed twice with water before being submitted for mass spectrometric analysis.

### Sample preparation for mass spectrometry analysis on RIPR-1::GFP Turbo-ID

The beads were resuspended in 50 µL 1 M urea and 50 mM ammonium bicarbonate. Disulfide bonds were reduced with 2 µL of 250 mM dithiothreitol (DTT) for 30 min at room temperature before adding 2 µL of 500 mM iodoacetamide and incubating for 30 min at room temperature in the dark. The remaining iodoacetamide was quenched with 1 µL of 250 mM DTT for 10 min. Proteins were digested with 150 ng trypsin (Trypsin Gold, Promega) in 1.5 µL 50 mM ammonium bicarbonate at room temperature for 90 minutes. The supernatant was transferred to a new tube. The beads were rinsed with 30 µL 1 M urea and 50 mM ammonium bicarbonate and pooled with the previous supernatant. Then the solution was digested further with another 150 ng trypsin (Trypsin Gold, Promega) in 1.5 µL 50 mM ammonium bicarbonate at 37°C overnight. The digest was stopped by the addition of trifluoroacetic acid (TFA) to a final concentration of 0.5%, and the peptides were desalted using C18 Stagetips^93^.

LC-MS analysis was performed on a Vanquish Neo UHPLC system (Thermo Scientific) coupled to an Orbitrap Exploris 480 mass spectrometer (Thermo Scientific). The system was equipped with a FAIMS pro interface (Thermo Scientific), a Nanospray Flex ion source (Thermo Scientific), coated emitter tips (PepSep, MSWil), and a Butterfly Portfolio Heater (Phoenix S&T).

Peptides were loaded onto a trap column (Acclaim PepMap 100 C18 HPLC Column, 5 mm × 300 µm, 5 μm particle size, Thermo Scientific) using 0.1% TFA as mobile phase, and separated on an analytical column (Acclaim PepMap 100 C18 HPLC Column, 50 cm × 75 µm, 2 μm particle size, Thermo Scientific), applying a linear gradient starting with a mobile phase of 98% solvent A (0.1% FA) and 2% solvent B (80% acetonitrile, 0.1% FA), increasing to 35% solvent B over 120 min at a flow rate of 230 nl/min. The analytical column was heated to 30°C.

The mass spectrometer was operated in data-dependent acquisition (DDA) mode with FAIMS compensation voltage (CV) set to alternate between −45 and −60, with 1.5 s cycle time per CV. Survey scans were acquired from 350-1500 m/z, normalized AGC target of 100%, resolution of 60,000. The most intense precursor ions (charge states +2 to +6) were selected for fragmentation using an isolation window of 1.4 m/z. Selected ions were analyzed with a maximum fill time of 50 ms, normalized AGC target of 100%, and resolution of 15,000 after HCD fragmentation with normalized collision energy of 30%. Monoisotopic precursor selection (MIPS) was set to “peptide” mode, the intensity threshold to 2.5E4, and selected precursors were dynamically excluded for 45 seconds with isotope exclusion enabled.

MS raw data were analyzed with FragPipe (23.1), using MSFragger (4.3)^94^, IonQuant (1.11.12)^95^, and Philosopher (5.1.2)^96^. The default FragPipe workflow for label free quantification (LFQ-MBR) was used, except “Normalize intensity across runs” was turned off. Cleavage specificity was set to Trypsin/P, with two missed cleavages allowed. The protein FDR was set to 1%. Carbamidomethyl was used as fixed cysteine modification; methionine oxidation, and protein N-terminal acetylation were specified as variable modifications. MS2 spectra were searched against the *Caenorhabditis elegans*1 protein per gene reference proteome from Uniprot (Proteome ID: UP000001940, release 2023_03), concatenated with a database of 379 common laboratory contaminants (release 2023.03, https://github.com/maxperutzlabs-ms/perutz-ms-contaminants).

Computational analysis was performed using Python and the in-house developed Python library MsReport (0.0.32)^97^. Only non-contaminant proteins identified with a minimum of two peptides and being quantified in at least one quantified replicate were considered for further analysis. LFQ protein intensities were log2-transformed and normalized across samples using the ModeNormalizer from MsReport. The ModeNormalizer method involves calculating log2 protein ratios for all pairs of samples and determining normalization factors based on the modes of all ratio distributions. Missing values were imputed with 16. iBAQ intensities were calculated by dividing protein intensities by the number of theoretically observable tryptic peptides between 6 and 30 amino acids. The Python library XlsxReport (0.1.1, https://github.com/hollenstein/xlsxreport) was used to create formatted Excel files summarizing the results of the proteomics experiments (Supplementary Table 3).

### Auxin treatment

All experiments involving exposure to auxin were carried out on NG medium plates containing 1 mM of 3-Indoleacetic acid (auxin) from a 440 mM stock dissolved in absolute Ethanol. Auxin-containing plates were seeded with 2x concentrated OP50 and kept protected from light at 4°C. Plates were used for no longer than three weeks. L4s or young adult animals were picked and maintained onto auxin plates for the indicated times. All strains employed in this study carry the *TIR1::mCherry* under the regulatory elements of *sun-1*, integrated on chromosome IV.

### DAPI staining

Synchronized animals from the relevant mutant strains and controls at the indicated age were dissected and processed as for normal immunofluorescence, except that blocking was omitted. After post-fixation in methanol, samples were washed three times for 5 minutes each in 1x PBST (1x PBS with 0.1% Tween) and 60 μl of DAPI in water (2 ng/ml) were applied on top of the samples. DAPI was allowed to stain for 1 minute and then slides were washed in the dark at room temperature for at least 20 minutes. Excess buffer was removed and a 15 μl drop of Vectashield was applied, before sealing the coverslips with nail polish.

### DNA substrates

All oligonucleotides used in the study were purchased from Eurofins Genomics and Sigma-Aldrich. For double-strand DNA substrates, equimolar amounts of individual oligonucleotides were annealed in hybridization buffer (50 mM Tris-HCl pH 7.5, 100 mM NaCl). The mixture was heated to 70 °C for 3 min and cooled slowly to room temperature.

For electrophoretic mobility shift assays (EMSA), ssDNA substrate (dT79, 5’ Cy3-labelled) and dsDNA substrate (90-mer, AAATAGACAGATCGCTGAGATAGGTGCCTCACTGATTAAGCATTGGTAACTGTCAGACCAAGTTTACTCATATATACTTTAGATTGATTT with its complementary 5’ FITC-labelled strand) were used.

For biolayer interferometry (BLI), ssDNA (dT80, 5’ biotinylated) and dsDNA (90-mer, AAATAGACAGATCGCTGAGATAGGTGCCTCACTGATTAAGCATTGGTAACTGTCAGACCAAGTTTACTCATATATACTTTAGATTGATTT with its complementary 5’ biotin-labelled strand) were used.

### Expression and purification of proteins

The codon-optimized *brc-2* gene was cloned into pCOLI_A_CBP or pLIB_CBP (#160155 or #160305, Addgene) using InteBac cloning strategy as previously described^98^. The coding sequence of *ripr-1* was PCR-amplified from in-house generated *C. elegans* cDNA and cloned into pLIB_12HIS (#160186, Addgene) using the same approach. For co-expression of BRC-2 and RIPR-1, a multicistronic construct was generated in the pBIG1a vector (#80611, Addgene) using the InteBac system. Baculovirus expression constructs were produced from pLIB plasmids using DH10Bac *E. coli* cells, and recombinant proteins were expressed in High Five insect cell. Insect cells expressing N-terminally His-tagged RIPR-1 were harvested and resuspended in the lysis buffer containing 50 mM Tris-HCl pH7.5, 100 mM KCl, 1 mM DTT, 0.1% NP-40, and protease inhibitors (2 µg/mL aprotinin, 5 µg/mL benzamidine, 10 µM chymostatin, 10 µM leupeptin, 1 µM pepstatin A and 1 mM phenylmethylsulfonyl fluoride). After sonication, the suspension was centrifuged at 15000 g for 45 minutes at 4 °C. Supernatant was mixed with Ni-NTA beads (Sigma) for 1 hour at 4 °C. Beads were subsequently washed with buffer containing 25 mM and 50 mM imidazole and proteins were eluted using buffer supplemented with 300 mM imidazole. For purification of CBP-tagged BRC-2/RIPR-1 complex, 2 mM CaCl2 was added to lysis buffer, and clarified lysates were incubated with calmodulin affinity resin for 30 minutes at 4 °C. The complex was eluted by buffer containing 10 mM EGTA without CaCl2. N-terminally CBP-tagged BRC-2 expressed in bacterial ArcticExpress cells was purified by the same procedure.

After affinity purification step, proteins and protein complex were concentrated using Vivaspin (10000 MWCO PES) and subjected to size-exclusion chromatography on Superdex 200 column equilibrated in buffer containing 50 mM Tris-HCl pH7.5, 100 mM NaCl and 0.01% NP40. Peak fractions were pooled and concentrated for subsequent biochemical analyses.

### Electrophoretic mobility shift assays (EMSA)

Purified RIPR-1, BRC-2, and the RIPR-1/BRC-2 complex were diluted in reaction buffer (50 mM Tris pH 7.5, 50 mM NaCl, 5 mM MgCl_2_). Proteins were incubated with 20 nM Cy3- or FITC-labelled DNA substrates in reaction buffer for 10 min at room temperature. Protein-DNA complexes were separated on 0.8% agarose gel in 0.5x TBE buffer at 70 V for 60 min at 4 °C. Gels were imaged by Typhoon™ laser-scanner (Cytiva), and band intensities were quantified by MultiGauge V3.2 software (Fujifilm). Obtained data were analysed using GraphPad Prism.

### Biolayer interferometry (BLI)

All BLI experiments were measured using a single-channel BLItz instrument (ForteBio) in Advanced Kinetics mode at room temperature with shaking at 2,200 rpm. The initial baseline was recorded for 30 sec in reaction buffer (50 mM Tris pH 7.5, 50 mM NaCl, 5 mM MgCl_2_, and 0.05% Tween-20). Biotinylated DNA substrates (15 nM ssDNA or dsDNA), were then loaded onto pre-hydrated streptavidin biosensors (SAX, Sartorius, Cat No. 18-5118) for 120 sec. After 30-second wash with reaction buffer, protein binding was monitored for 180 sec in reaction buffer. Protein dissociation from DNA substrate was monitored for an additional 180 sec in the same buffer but including 100 mM NaCl. Data were analysed using single exponential fit in BLItz Pro 1.2 software using global fitting mode.

## Data availability

All data generated or analyzed in this study are included in this published article (and its Supplementary information files).

## ACKNOWLEDGMENTS

We would like to thank Simone Köhler for sharing the *V5::rad-51b* tagged line prior publication and Batool Ossareh-Nazari and Lionel Pintard for sharing the protocol for the TurboID sample preparation. We acknowledge CEITEC Proteomics Core Facility of CIISB, Instruct-CZ Centre, supported by MEYS CR (LM2023042, CZ.02.01.01/00/23_015/0008175, e-INFRA CZ (ID:90254)) for the analysis performed on FLAG::BRC-2 pull downs. We acknowledge the core facility CELLIM supported by the Czech-BioImaging large RI project (LM2023050 funded by MEYS CR) for their support with obtaining scientific data presented in this paper. Proteomics analyses on Turbo-ID samples were performed by the Mass Spectrometry Facility at Max Perutz Labs using the VBCF instrument pool. GFP::MSH-5 images were acquired at the Max Perutz Labs’ BioOptics facility and we acknowledge their BioOptics team for training and support in image acquisition. Some strains were provided by the CGC, which is funded by NIH Office of Research Infrastructure Programs (P40 OD010440). The Silva laboratory is funded by the Czech Science Foundation (GA26-20768S), the Krejci laboratory is funded by the Czech Science Foundation (21-22593X) and European Union’s Horizon Europe Research and Innovation Programme (101158508), the Jantsch laboratory is funded by the Austrian Science Fund (FWF) grant PAT2512023 (Grant-DOI 10.55776/PAT2512023) and the SFB project F 8805-B (Grant-DOI 10.55776/F88).

## Author Contributions

NS acquired funding, designed the research and performed most of the experiments with the technical support of JB and NBAB; MS and ML performed BRC-2-RIPR-1 expression, purification and all the *in vitro* assays; SSG acquired images of GFP::MSH-5 and produced extracts and IPs for TurboID experiments; NS, VJ and LK provided supervision; NS wrote the manuscript, with editing inputs from LK.

## Competing Interests

The authors declare no competing interests.

## Notes

### Competing Interest Statement

The authors have declared no competing interest.

## REFERENCES

1. Keeney, S., Giroux, C. N. & Kleckner, N. Meiosis-specific DNA double-strand breaks are catalyzed by Spo11, a member of a widely conserved protein family. Cell 88, 375–384 (1997).

2. Gray, S. & Cohen, P. E. Control of Meiotic Crossovers: From Double-Strand Break Formation to Designation. Annu. Rev. Genet. 50, 175–210 (2016).

3. Zickler, D. & Kleckner, N. Meiosis: Dances Between Homologs. Annu. Rev. Genet. 57, 1–63 (2023).

4. Dernburg, A. F. et al. Meiotic Recombination in C. elegans Initiates by a Conserved Mechanism and Is Dispensable for Homologous Chromosome Synapsis. Cell 94, 387–398 (1998).

5. Ito, M., Fujita, Y. & Shinohara, A. Positive and negative regulators of RAD51/DMC1 in homologous recombination and DNA replication. DNA Repair 134, 103613 (2024).

6. Borde, V. The multiple roles of the Mre11 complex for meiotic recombination. Chromosome Res 15, 551–563 (2007).

7. Sharan, S. K. et al. Embryonic lethality and radiation hypersensitivity mediated by Rad51 in mice lacking Brca2. Nature 386, 804–810 (1997).

8. Davies, A. A. et al. Role of BRCA2 in Control of the RAD51 Recombination and DNA Repair Protein. Molecular Cell 7, 273–282 (2001).

9. Wong, A. K. C., Pero, R., Ormonde, P. A., Tavtigian, S. V. & Bartel, P. L. RAD51 Interacts with the Evolutionarily Conserved BRC Motifs in the Human Breast Cancer Susceptibility Gene brca2. Journal of Biological Chemistry 272, 31941–31944 (1997).

10. Dray, E., Siaud, N., Dubois, E. & Doutriaux, M.-P. Interaction between Arabidopsis Brca2 and Its Partners Rad51, Dmc1, and Dss1. Plant Physiology 140, 1059–1069 (2006).

11. Jensen, R. B., Carreira, A. & Kowalczykowski, S. C. Purified human BRCA2 stimulates RAD51-mediated recombination. Nature 467, 678–683 (2010).

12. Thorslund, T., Esashi, F. & West, S. C. Interactions between human BRCA2 protein and the meiosis-specific recombinase DMC1. EMBO J 26, 2915–2922 (2007).

13. Siaud, N. et al. Brca2 is involved in meiosis in Arabidopsis thaliana as suggested by its interaction with Dmc1. EMBO J 23, 1392–1401 (2004).

14. Wooster, R. et al. Identification of the breast cancer susceptibility gene BRCA2. Nature 378, 789–792 (1995).

15. Yu, V. P. C. C. et al. Gross chromosomal rearrangements and genetic exchange between nonhomologous chromosomes following BRCA2 inactivation. Genes Dev. 14, 1400–1406 (2000).

16. Venkitaraman, A. R. Cancer Susceptibility and the Functions of BRCA1 and BRCA2. Cell 108, 171–182 (2002).

17. King, M.-C., Marks, J. H. & Mandell, J. B. Breast and Ovarian Cancer Risks Due to Inherited Mutations in *BRCA1* and *BRCA2*. Science 302, 643–646 (2003).

18. Xia, B. et al. Control of BRCA2 Cellular and Clinical Functions by a Nuclear Partner, PALB2. Molecular Cell 22, 719–729 (2006).

19. Takemoto, K. et al. Meiosis-Specific C19orf57/4930432K21Rik/BRME1 Modulates Localization of RAD51 and DMC1 to DSBs in Mouse Meiotic Recombination. Cell Reports 31, 107686 (2020).

20. Zhang, J. et al. The BRCA2-MEILB2-BRME1 complex governs meiotic recombination and impairs the mitotic BRCA2-RAD51 function in cancer cells. Nat Commun 11, 2055 (2020).

21. Simhadri, S. et al. Male Fertility Defect Associated with Disrupted BRCA1-PALB2 Interaction in Mice. Journal of Biological Chemistry 289, 24617–24629 (2014).

22. Martin, J. S., Winkelmann, N., Petalcorin, M. I. R., McIlwraith, M. J. & Boulton, S. J. RAD-51-Dependent and -Independent Roles of a *Caenorhabditis elegans* BRCA2-Related Protein during DNA Double-Strand Break Repair. Mol Cell Biol 25, 3127–3139 (2005).

23. Alpi, A., Pasierbek, P., Gartner, A. & Loidl, J. Genetic and cytological characterization of the recombination protein RAD-51 in Caenorhabditis elegans. Chromosoma 112, 6–16 (2003).

24. Colaiácovo, M. P. et al. Synaptonemal Complex Assembly in C. elegans Is Dispensable for Loading Strand-Exchange Proteins but Critical for Proper Completion of Recombination. Developmental Cell 5, 463–474 (2003).

25. Kelly, K. O., Dernburg, A. F., Stanfield, G. M. & Villeneuve, A. M. Caenorhabditis elegans msh-5 is required for both normal and radiation-induced meiotic crossing over but not for completion of meiosis. Genetics 156, 617–630 (2000).

26. Yokoo, R. et al. COSA-1 Reveals Robust Homeostasis and Separable Licensing and Reinforcement Steps Governing Meiotic Crossovers. Cell 149, 75–87 (2012).

27. Rinaldo, C., Bazzicalupo, P., Ederle, S., Hilliard, M. & La Volpe, A. Roles for *Caenorhabditis elegans rad-51* in Meiosis and in Resistance to Ionizing Radiation During Development. Genetics 160, 471–479 (2002).

28. Trivedi, S., Blazícková, J. & Silva, N. PARG and BRCA1–BARD1 cooperative function regulates DNA repair pathway choice during gametogenesis. Nucleic Acids Research 50, 12291–12308 (2022).

29. Bianco, P. R. OB-fold Families of Genome Guardians: A Universal Theme Constructed From the Small β-barrel Building Block. Front. Mol. Biosci. 9, 784451 (2022).

30. Flynn, R. L. & Zou, L. Oligonucleotide/oligosaccharide-binding fold proteins: a growing family of genome guardians. Critical Reviews in Biochemistry and Molecular Biology 45, 266–275 (2010).

31. Woglar, A. et al. Matefin/SUN-1 Phosphorylation Is Part of a Surveillance Mechanism to Coordinate Chromosome Synapsis and Recombination with Meiotic Progression and Chromosome Movement. PLoS Genet 9, e1003335 (2013).

32. Goodyer, W. et al. HTP-3 Links DSB Formation with Homolog Pairing and Crossing Over during C. elegans Meiosis. Developmental Cell 14, 263–274 (2008).

33. Severson, A. F., Ling, L., van Zuylen, V. & Meyer, B. J. The axial element protein HTP-3 promotes cohesin loading and meiotic axis assembly in C. elegans to implement the meiotic program of chromosome segregation. Genes & Development 23, 1763–1778 (2009).

34. Couteau, F. HTP-1 coordinates synaptonemal complex assembly with homolog alignment during meiosis in C. elegans. Genes & Development 19, 2744–2756 (2005).

35. Martinez-Perez, E. HTP-1-dependent constraints coordinate homolog pairing and synapsis and promote chiasma formation during C. elegans meiosis. Genes & Development 19, 2727–2743 (2005).

36. Couteau, F., Nabeshima, K., Villeneuve, A. & Zetka, M. A Component of C. elegans Meiotic Chromosome Axes at the Interface of Homolog Alignment, Synapsis, Nuclear Reorganization, and Recombination. Current Biology 14, 585–592 (2004).

37. Severson, A. F. & Meyer, B. J. Divergent kleisin subunits of cohesin specify mechanisms to tether and release meiotic chromosomes. eLife 3, e03467 (2014).

38. MacQueen, A. J. Synapsis-dependent and -independent mechanisms stabilize homolog pairing during meiotic prophase in C. elegans. Genes & Development 16, 2428–2442 (2002).

39. Smolikov, S. et al. SYP-3 Restricts Synaptonemal Complex Assembly to Bridge Paired Chromosome Axes During Meiosis in *Caenorhabditis elegans*. Genetics 176, 2015–2025 (2007).

40. Smolikov, S., Schild-Prüfert, K. & Colaiácovo, M. P. A Yeast Two-Hybrid Screen for SYP-3 Interactors Identifies SYP-4, a Component Required for Synaptonemal Complex Assembly and Chiasma Formation in Caenorhabditis elegans Meiosis. PLoS Genet 5, e1000669 (2009).

41. Blundon, J. M. et al. Skp1 proteins are structural components of the synaptonemal complex in *C. elegans*. Sci. Adv. 10, eadl4876 (2024).

42. Hurlock, M. E. et al. Identification of novel synaptonemal complex components in C. elegans. Journal of Cell Biology 219, e201910043 (2020).

43. Blazickova, J. et al. Overlapping and separable activities of BRA-2 and HIM-17 promote occurrence and regulation of pairing and synapsis during Caenorhabditis elegans meiosis. Nat Commun 16, 2516 (2025).

44. Machovina, T. S. et al. A Surveillance System Ensures Crossover Formation in C. elegans. Current Biology 26, 2873–2884 (2016).

45. Pattabiraman, D., Roelens, B., Woglar, A. & Villeneuve, A. M. Meiotic recombination modulates the structure and dynamics of the synaptonemal complex during C. elegans meiosis. PLoS Genet 13, e1006670 (2017).

46. Yin, Y. & Smolikove, S. Impaired Resection of Meiotic Double-Strand Breaks Channels Repair to Nonhomologous End Joining in Caenorhabditis elegans. Mol Cell Biol 33, 2732–2747 (2013).

47. Lemmens, B. B. L. G., Johnson, N. M. & Tijsterman, M. COM-1 Promotes Homologous Recombination during Caenorhabditis elegans Meiosis by Antagonizing Ku-Mediated Non-Homologous End Joining. PLoS Genet 9, e1003276 (2013).

48. Girard, C., Roelens, B., Zawadzki, K. A. & Villeneuve, A. M. Interdependent and separable functions of *Caenorhabditis elegans* MRN-C complex members couple formation and repair of meiotic DSBs. Proc Natl Acad Sci USA 115, E4443–E4452 (2018).

49. Koury, E., Harrell, K. & Smolikove, S. Differential RPA-1 and RAD-51 recruitment in vivo throughout the C. elegans germline, as revealed by laser microirradiation. Nucleic Acids Research 46, 748–764 (2018).

50. Gumienny, T. L., Lambie, E., Hartwieg, E., Horvitz, H. R. & Hengartner, M. O. Genetic control of programmed cell death in the Caenorhabditis elegans hermaphrodite germline. Development 126, 1011–1022 (1999).

51. Bhalla, N. & Dernburg, A. F. A Conserved Checkpoint Monitors Meiotic Chromosome Synapsis in *Caenorhabditis elegans*. Science 310, 1683–1686 (2005).

52. Schumacher, B., Hofmann, K., Boulton, S. & Gartner, A. The C. elegans homolog of the p53 tumor suppressor is required for DNA damage-induced apoptosis. Current Biology 11, 1722–1727 (2001).

53. Zhang, L., Ward, J. D., Cheng, Z. & Dernburg, A. F. The auxin-inducible degradation (AID) system enables versatile conditional protein depletion in *C. elegans*. Development dev.129635 (2015) doi:10.1242/dev.129635.

54. Hicks, T. et al. Continuous double-strand break induction and their differential processing sustain chiasma formation during Caenorhabditis elegans meiosis. Cell Reports 40, 111403 (2022).

55. Chin, G. M. C. elegans mre-11 is required for meiotic recombination and DNA repair but is dispensable for the meiotic G2 DNA damage checkpoint. Genes & Development 15, 522–534 (2001).

56. Hayashi, M., Chin, G. M. & Villeneuve, A. M. C. elegans germ cells switch between distinct modes of double-strand break repair during meiotic prophase progression. PLoS Genet 3, e191 (2007).

57. Harrell, K., Day, M. & Smolikove, S. Recruitment of MRE-11 to complex DNA damage is modulated by meiosis-specific chromosome organization. Mutat Res 822, 111743 (2021).

58. Hefel, A. et al. RPA complexes in *Caenorhabditis elegans* meiosis; unique roles in replication, meiotic recombination and apoptosis. Nucleic Acids Research 49, 2005–2026 (2021).

59. Neves, A. R. R., Čavka, I., Rausch, T. & Köhler, S. Crossovers are regulated by a conserved and disordered synaptonemal complex domain. Nucleic Acids Research 53, gkaf095 (2025).

60. Adamo, A. et al. BRC-1 acts in the inter-sister pathway of meiotic double-strand break repair. EMBO Rep 9, 287–292 (2008).

61. Janisiw, E., Dello Stritto, M. R., Jantsch, V. & Silva, N. BRCA1-BARD1 associate with the synaptonemal complex and pro-crossover factors and influence RAD-51 dynamics during Caenorhabditis elegans meiosis. PLoS Genet 14, e1007653 (2018).

62. Li, Q. et al. The tumor suppressor BRCA1-BARD1 complex localizes to the synaptonemal complex and regulates recombination under meiotic dysfunction in Caenorhabditis elegans. PLoS Genet 14, e1007701 (2018).

63. Toraason, E. et al. BRCA1/BRC-1 and SMC-5/6 regulate DNA repair pathway engagement during Caenorhabditis elegans meiosis. eLife 13, e80687 (2024).

64. Muller, H. J. The Mechanism of Crossing-Over. The American Naturalist 50, 193–221 (1916).

65. Zalevsky, J., MacQueen, A. J., Duffy, J. B., Kemphues, K. J. & Villeneuve, A. M. Crossing over during Caenorhabditis elegans meiosis requires a conserved MutS-based pathway that is partially dispensable in budding yeast. Genetics 153, 1271–1283 (1999).

66. Bhalla, N., Wynne, D. J., Jantsch, V. & Dernburg, A. F. ZHP-3 Acts at Crossovers to Couple Meiotic Recombination with Synaptonemal Complex Disassembly and Bivalent Formation in C. elegans. PLoS Genet 4, e1000235 (2008).

67. Jagut, M. et al. Separable Roles for a Caenorhabditis elegans RMI1 Homolog in Promoting and Antagonizing Meiotic Crossovers Ensure Faithful Chromosome Inheritance. PLoS Biol 14, e1002412 (2016).

68. Wicky, C. et al. Multiple Genetic Pathways Involving the *Caenorhabditis elegans* Bloom’s Syndrome Genes *him-6*, *rad-51*, and *top-3* Are Needed To Maintain Genome Stability in the Germ Line. Molecular and Cellular Biology 24, 5016–5027 (2004).

69. Engebrecht, J. et al. Loss of meiotic double strand breaks triggers recruitment of recombination-independent pro-crossover factors in C. elegans spermatogenesis. PLoS Genet 21, e1011763 (2025).

70. Holzer, E., Rumpf-Kienzl, C., Falk, S. & Dammermann, A. A modified TurboID approach identifies tissue-specific centriolar components in C. elegans. PLoS Genet 18, e1010150 (2022).

71. Petalcorin, M. I. R., Sandall, J., Wigley, D. B. & Boulton, S. J. CeBRC-2 Stimulates D-loop Formation by RAD-51 and Promotes DNA Single-strand Annealing. Journal of Molecular Biology 361, 231–242 (2006).

72. Pellegrini, L. et al. Insights into DNA recombination from the structure of a RAD51–BRCA2 complex. Nature 420, 287–293 (2002).

73. Park, J.-Y., Zhang, F. & Andreassen, P. R. PALB2: The hub of a network of tumor suppressors involved in DNA damage responses. Biochimica et Biophysica Acta (BBA) - Reviews on Cancer 1846, 263–275 (2014).

74. The Breast Cancer Susceptibility Collaboration (UK) et al. PALB2, which encodes a BRCA2-interacting protein, is a breast cancer susceptibility gene. Nat Genet 39, 165–167 (2007).

75. Tischkowitz, M. et al. Analysis of *PALB2* / *FANCN* -associated breast cancer families. Proc. Natl. Acad. Sci. U.S.A. 104, 6788–6793 (2007).

76. Amir, Mohd., et al. A Systems View of the Genome Guardians: Mapping the Signaling Circuitry Underlying Oligonucleotide/Oligosaccharide-Binding Fold Proteins. OMICS: A Journal of Integrative Biology 24, 518–530 (2020).

77. Nguyen, D.-D., Kim, E. Y., Sang, P. B. & Chai, W. Roles of OB-Fold Proteins in Replication Stress. Front. Cell Dev. Biol. 8, 574466 (2020).

78. Yang, H. et al. BRCA2 Function in DNA Binding and Recombination from a BRCA2-DSS1-ssDNA Structure. Science 297, 1837–1848 (2002).

79. Shivji, M. K. K. et al. The BRC repeats of human BRCA2 differentially regulate RAD51 binding on single- versus double-stranded DNA to stimulate strand exchange. Proc. Natl. Acad. Sci. U.S.A. 106, 13254–13259 (2009).

80. Carreira, A. et al. The BRC Repeats of BRCA2 Modulate the DNA-Binding Selectivity of RAD51. Cell 136, 1032–1043 (2009).

81. Zhao, W. et al. Promotion of BRCA2-Dependent Homologous Recombination by DSS1 via RPA Targeting and DNA Mimicry. Molecular Cell 59, 176–187 (2015).

82. Bell, J. C., Dombrowski, C. C., Plank, J. L., Jensen, R. B. & Kowalczykowski, S. C. BRCA2 chaperones RAD51 to single molecules of RPA-coated ssDNA. Proc. Natl. Acad. Sci. U.S.A. 120, e2221971120 (2023).

83. Liu, J., Doty, T., Gibson, B. & Heyer, W.-D. Human BRCA2 protein promotes RAD51 filament formation on RPA-covered single-stranded DNA. Nat Struct Mol Biol 17, 1260–1262 (2010).

84. Ding, J. et al. ssDNA accessibility of Rad51 is regulated by orchestrating multiple RPA dynamics. Nat Commun 14, 3864 (2023).

85. Ma, C. J., Gibb, B., Kwon, Y., Sung, P. & Greene, E. C. Protein dynamics of human RPA and RAD51 on ssDNA during assembly and disassembly of the RAD51 filament. Nucleic Acids Res 45, 749–761 (2017).

86. Pokhrel, N. et al. Dynamics and selective remodeling of the DNA-binding domains of RPA. Nat Struct Mol Biol 26, 129–136 (2019).

87. Strand, L. G. et al. Active maintenance of meiosis-specific chromosome structures in *Caenorhabditis elegans* by the deubiquitinase DUO-1. Proc. Natl. Acad. Sci. U.S.A. 123, e2532671123 (2026).

88. Libuda, D. E., Uzawa, S., Meyer, B. J. & Villeneuve, A. M. Meiotic chromosome structures constrain and respond to designation of crossover sites. Nature 502, 703–706 (2013).

89. Cahoon, C. K., Helm, J. M. & Libuda, D. E. Synaptonemal Complex Central Region Proteins Promote Localization of Pro-crossover Factors to Recombination Events During *Caenorhabditis elegans* Meiosis. Genetics 213, 395–409 (2019).

90. Huang, Y. et al. DSS1 restrains BRCA2’s engagement with dsDNA for homologous recombination, replication fork protection, and R-loop homeostasis. Nat Commun 15, 7081 (2024).

91. Silva, N. et al. The Fidelity of Synaptonemal Complex Assembly Is Regulated by a Signaling Mechanism that Controls Early Meiotic Progression. Developmental Cell 31, 503–511 (2014).

92. Stejskal, K., Potěšil, D. & Zdráhal, Z. Suppression of Peptide sample losses in autosampler vials. J. Proteome Res. 12, 3057–3062 (2013).

93. Rappsilber, J., Mann, M. & Ishihama, Y. Protocol for micro-purification, enrichment, pre-fractionation and storage of peptides for proteomics using StageTips. Nat Protoc 2, 1896–1906 (2007).

94. Kong, A. T., Leprevost, F. V., Avtonomov, D. M., Mellacheruvu, D. & Nesvizhskii, A. I. MSFragger: ultrafast and comprehensive peptide identification in mass spectrometry–based proteomics. Nat Methods 14, 513–520 (2017).

95. Yu, F., Haynes, S. E. & Nesvizhskii, A. I. IonQuant Enables Accurate and Sensitive Label-Free Quantification With FDR-Controlled Match-Between-Runs. Molecular & Cellular Proteomics 20, 100077 (2021).

96. Da Veiga Leprevost, F., et al. Philosopher: a versatile toolkit for shotgun proteomics data analysis. Nat Methods 17, 869–870 (2020).

97. Hollenstein, D. M. et al. Chemical Acetylation of Ligands and Two-Step Digestion Protocol for Reducing Codigestion in Affinity Purification–Mass Spectrometry. J. Proteome Res. 22, 3383–3391 (2023).

98. Altmannova, V., Blaha, A., Astrinidis, S., Reichle, H. & Weir, J. R. INTEBAC : An integrated bacterial and baculovirus expression vector suite. Protein Science 30, 108–114 (2021).

