## Supplementary Figures for "BRC-2/BRCA2-RIPR-1 mediated constraints on homologous recombination execution are spatiotemporally regulated during meiosis"

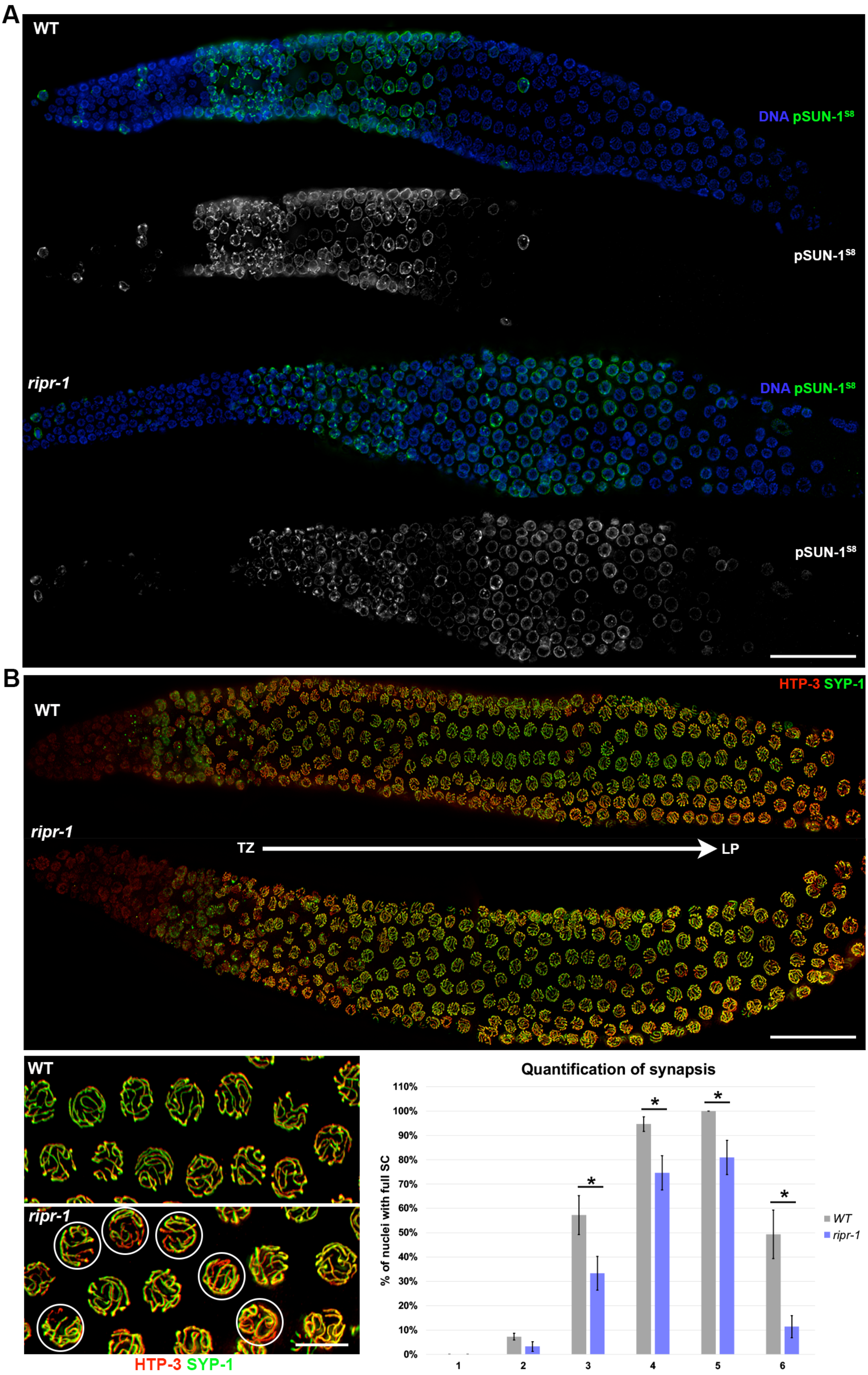
A
WT
DNA pSUN-1S8
pSUN-1S8
ripr-1
DNA pSUN-1S8
pSUN-1S8
B
HTP-3 SYP-1
WT
ripr-1
TZ
LP
WT
ripr-1
HTP-3 SYP-1
Quantification of synapsis
% of nuclei with full SC
110%
100%
90%
80%
70%
60%
50%
40%
30%
20%
10%
0%
1
2
3
4
5
6
*
*
*
*
WT
ripr-1

**Supp. Fig. 1. Loss of *ripr-1* prolongs CHK-2-mediated signaling and perturbs SC assembly.**

**(A)** Whole-mount gonads from WT and *ripr-1* mutant animals stained for phosphoSUN-1$^{S8}$ (green) and counterstained by DAPI (blue). Scale bar 20 μm. **(B)** Top: whole-mount gonads from WT and *ripr-1* mutant animals stained for HTP-3 (red) and SYP-1 (green). White arrow shows orientation of nuclei from transition zone (TZ) toward Late Pachytene (LP). Scale bar 20 μm. Bottom: representative images of magnified LP nuclei from WT controls and *ripr-1* mutants stained for HTP-3/SYP-1. White circles highlight nuclei with incomplete synapsis as observed by chromosomal regions stained by HTP-3 but not SYP-1. Scale bar 5 μm. On the right side, chart showing quantification of SC assembly in WT and *ripr-1* mutant worms across the germ line. Bars indicate S.E.M. and asterisks denote statistical significance as assessed by $X^2$ test (P=0.05)

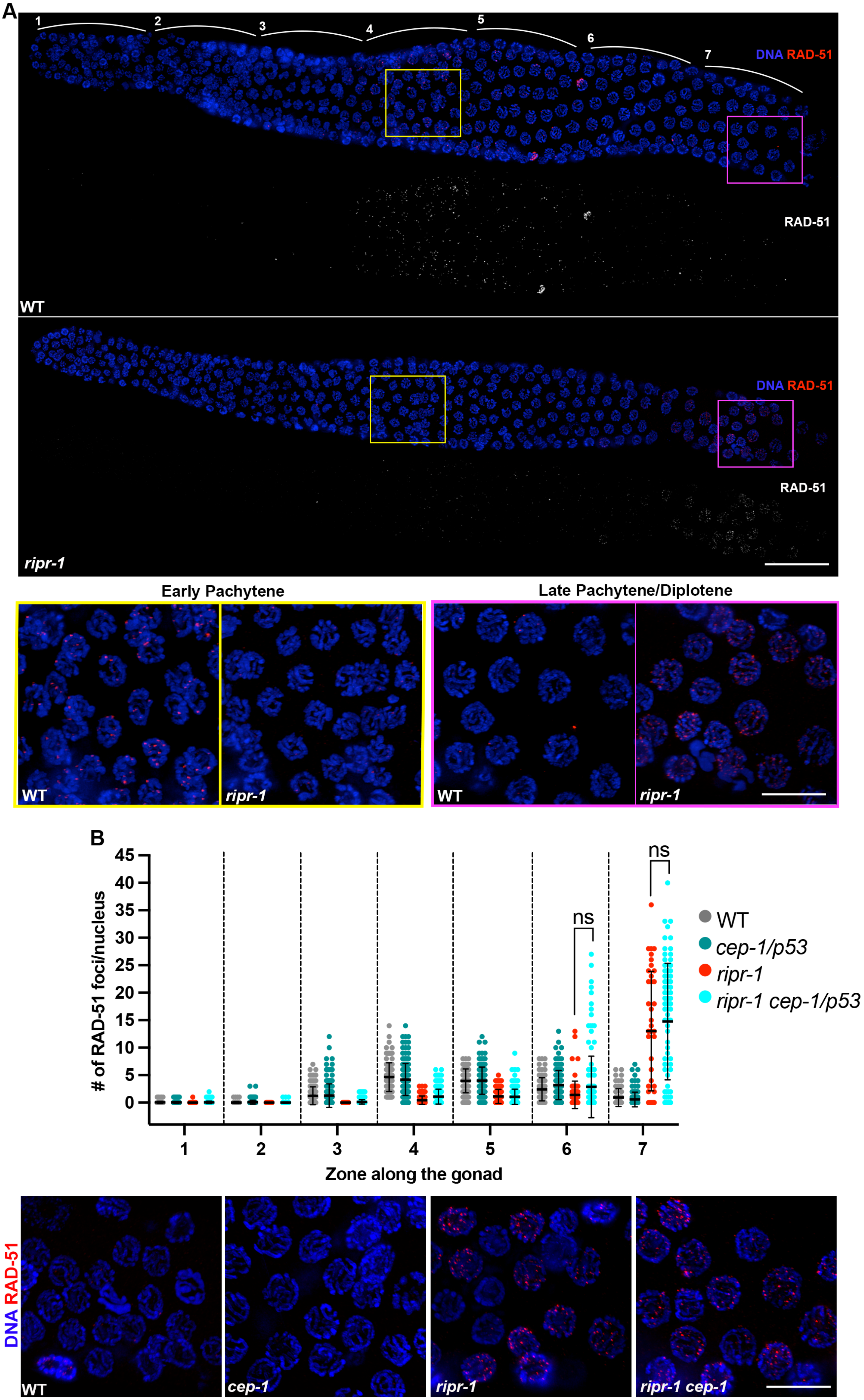
A
1
2
3
4
5
6
7
DNA RAD-51
RAD-51
WT
DNA RAD-51
RAD-51
ripr-1
Early Pachytene
Late Pachytene/Diplotene
WT
ripr-1
WT
ripr-1
B
### of RAD-51 foci/nucleus
Zone along the gonad
ns
ns
WT
cep-1/p53
ripr-1
ripr-1 cep-1/p53
DNA RAD-51
WT
cep-1
ripr-1
ripr-1 cep-1

**Supp. Fig. 2. Removal of RIPR-1 results in dramatic RAD-51 accumulation in late pachytene nuclei that does not depend on apoptosis.**

**(A)** Whole-mount germ lines stained with RAD-51 antibodies and counterstained by DAPI in WT and *ripr-1* mutant germ lines. Zoning adopted for foci quantification is shown and colored square highlight the regions of the gonad magnified underneath. Scale bar 10 μm. **(B)** Top: quantification of RAD-51 foci along the germ line in the indicated genetic backgrounds. Bars indicate average with S.D. and *ns*=not significant (T test). Bottom: representative images of late pachytene cells from the indicated mutant and control animals, immunoassayed for RAD-51 (red) and DAPI (blue). Scale bar 10 μm.

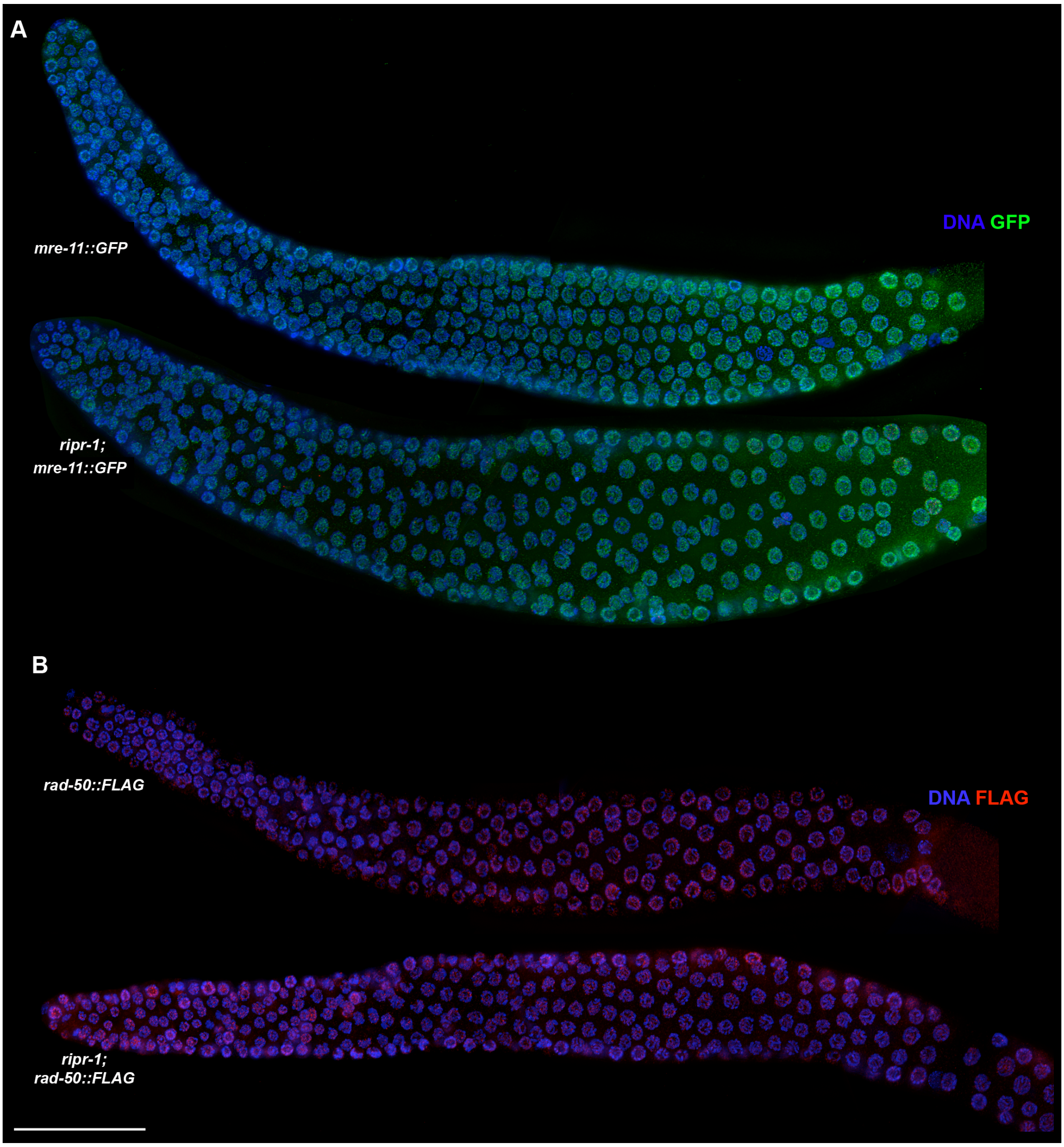


**Supp. Fig. 3. RIPR-1 is dispensable for localization of MRE-11 and RAD-50.**

**(A)** Whole-mount germlines immunoassayed for GFP (MRE-11) and **(B)** FLAG (RAD-50) in control animals and *ripr-1* mutants**.** Scale bar 20 μm.

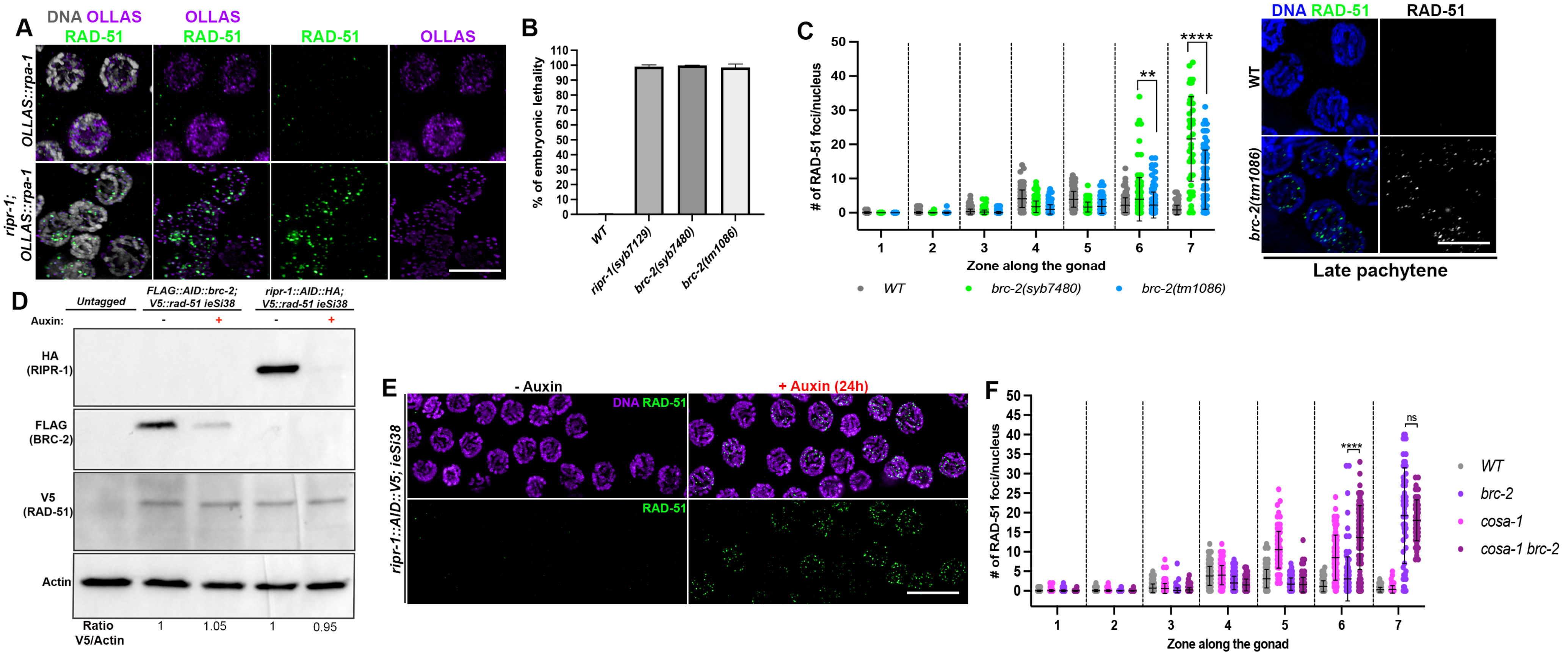


**Supp. Fig. 4. RAD-51 similarly accumulates in *ripr-1* and different *brc-2* mutant alleles.**

**(A)** Late pachytene cells immunoassayed for OLLAS::RPA-1 (purple) and RAD-51 (green) in the indicated genotypes. Scale bar 5 μm. **(B)** Assessment of viability levels in the indicated mutant backgrounds and WT controls. Bars indicate average of embryonic lethality and S.D. **(C)** Left: quantification of RAD-51 foci in the indicated genotypes. Bars represent average with S.D. Asterisks denote statistical significance assessed by T test (***p*=0.0068, *****p<0.0001*). Right: representative images of late pachytene nuclei stained for RAD-51 (green) and counterstained by

DAPI (blue). Scale bar 5 μm. **(D)** Western blot analysis on whole cell extracts from the indicated genetic backgrounds and exposure conditions to auxin. **(E)** Representative images of late pachytene nuclei immunoassayed for RAD-51 (green) and counterstained by DAPI (purple) in the indicated genetic background before and after 24h exposure to auxin. Scale bar 5 μm. **(F)** Quantification of RAD-51 foci in the indicated genotypes. Bars represent average with S.D. Asterisks denote statistical significance assessed by T test (***p*=0.0068, *****p<0.0001, ns*=non-significant).

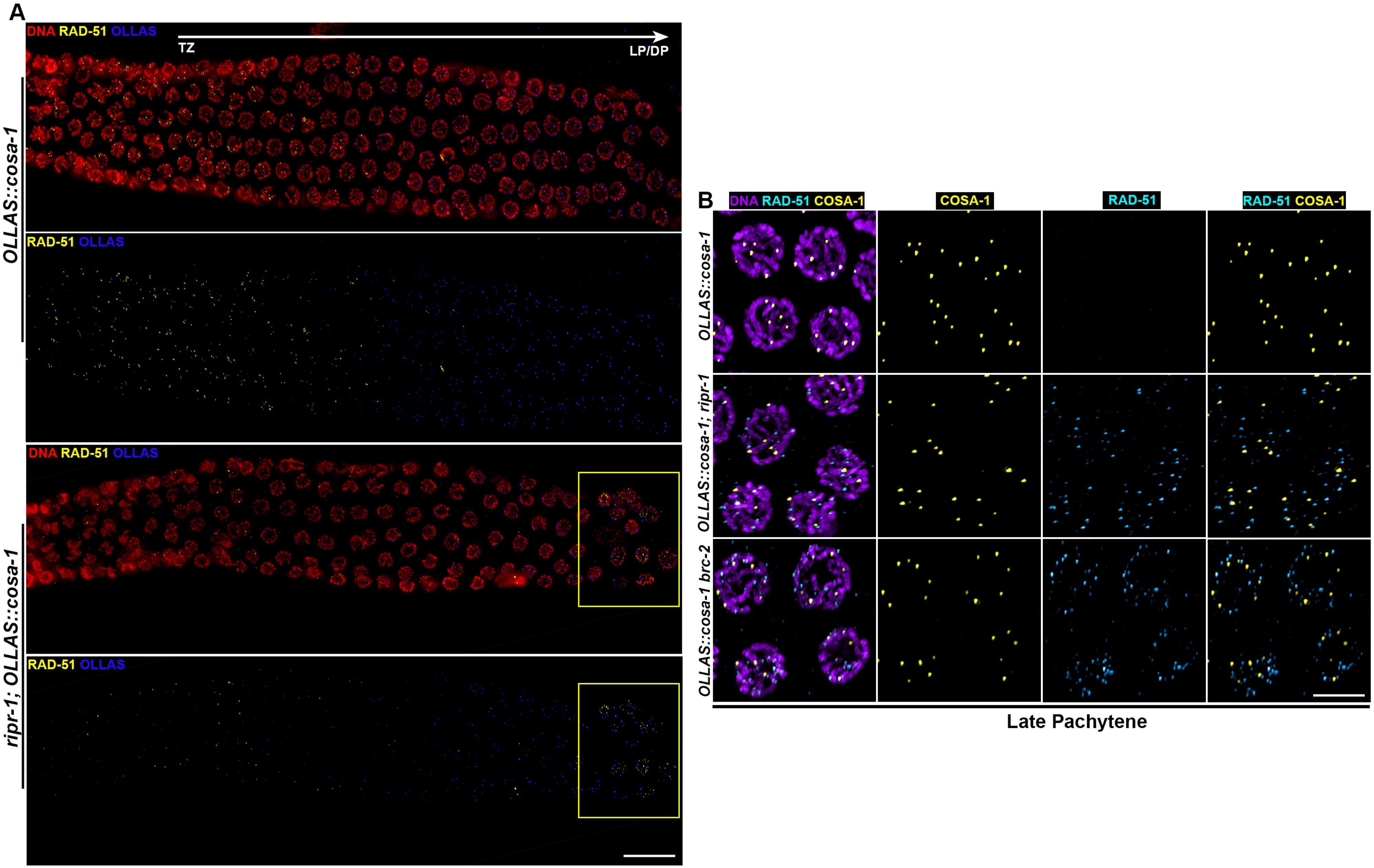
A
DNA RAD-51 OLLAS
TZ
LP/DP
OLLAS::cosa-1
RAD-51 OLLAS
DNA RAD-51 OLLAS
ripr-1; OLLAS::cosa-1
RAD-51 OLLAS
B
DNA RAD-51 COSA-1
COSA-1
RAD-51
RAD-51 COSA-1
OLLAS::cosa-1
OLLAS::cosa-1; ripr-1
OLLAS::cosa-1 brc-2
Late Pachytene

**Supp. Fig. 5. Loss of RIPR-1-BRC-2 triggers simultaneous recruitment of COSA-1 and RAD-51 in late pachytene cells.**

**(A)** Representative images of partial germ lines from the indicated genetic backgrounds, immunoassayed for OLLAS::COSA-1 (blue), RAD-51 (yellow) and counterstained by DAPI (red). Scale bar 5 μm. **(B)** Late pachytene cells from the indicated genotypes, immunoassayed for OLLAS::COSA-1 (yellow) and RAD-51 (cyan), counterstained by DAPI (purple). Scale bar 5 μm.

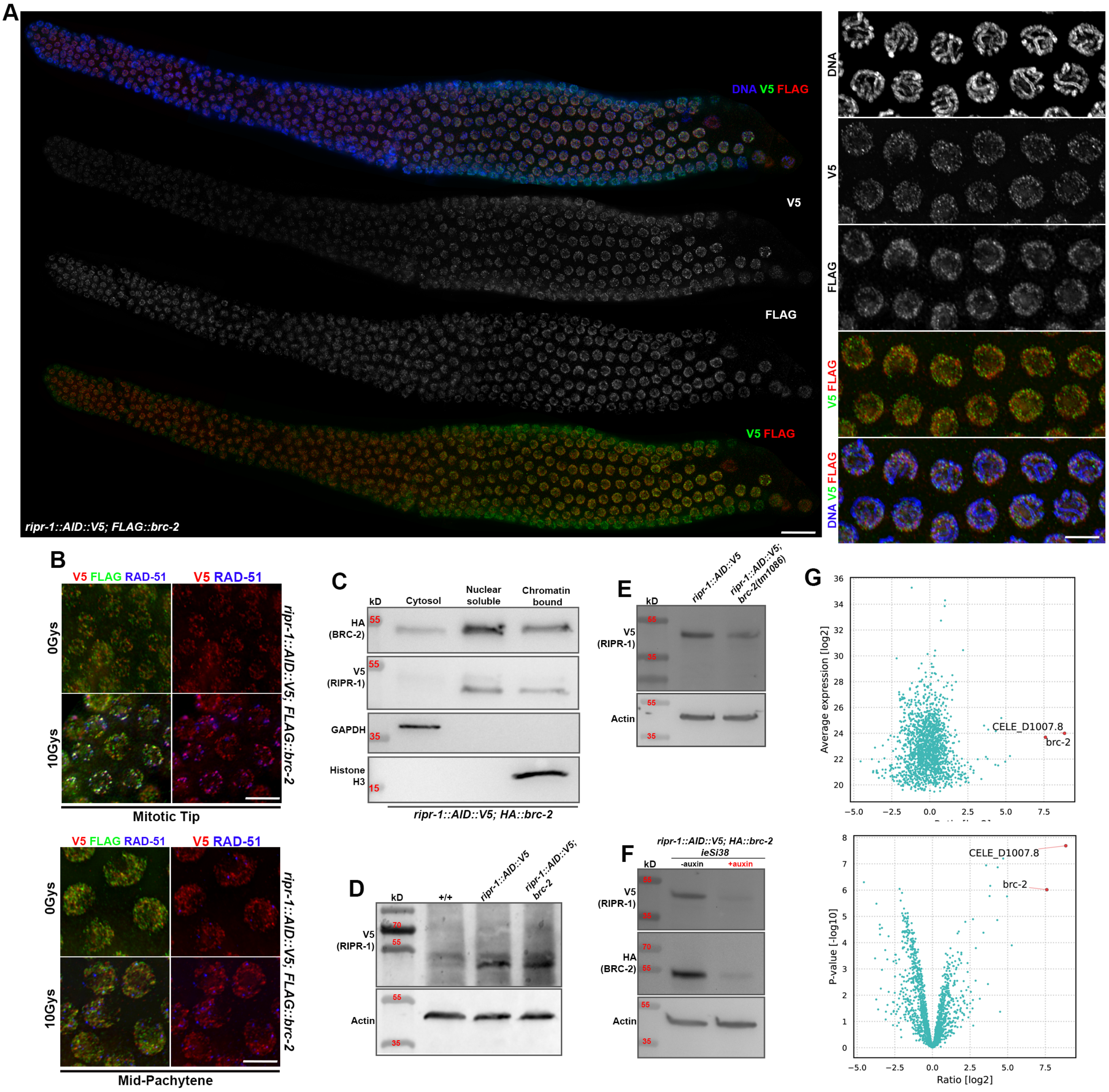


**Supp. Fig. 6. RIPR-1 and BRC-2 loading is mutually dependent.**

**(A)** Left: whole-mount gonads from *ripr-1::AID::V5; FLAG::brc-2* worms immunostained with V5 (RIPR-1) and FLAG (BRC-2) antibodies. Right: magnification of mid-pachytene cells. Scale bar 3 μm. **(B)** Representative images of nuclei from the mitotic tip (top) and mid-pachytene stage (bottom) of the indicated genetic background and exposure to IR, immunoassayed with V5 (RIPR-1, red), FLAG (BRC-2, green) and RAD-51 (blue) antibodies. Scale bar 3 μm. **(C)** Western blot analysis on fractionated protein extracts from *ripr-1::AID::V5; HA::brc-2* worms and probed with the indicated antibodies. **(D-F)** Western blot analysis on whole-cell extracts on the indicated genotypes and exposure conditions to auxin, probed with the specified antibodies. **(G)** Volcano plots showing expression and enrichment of RIPR-1 (D1007.8) and BRC-2 in TurboID experiments.

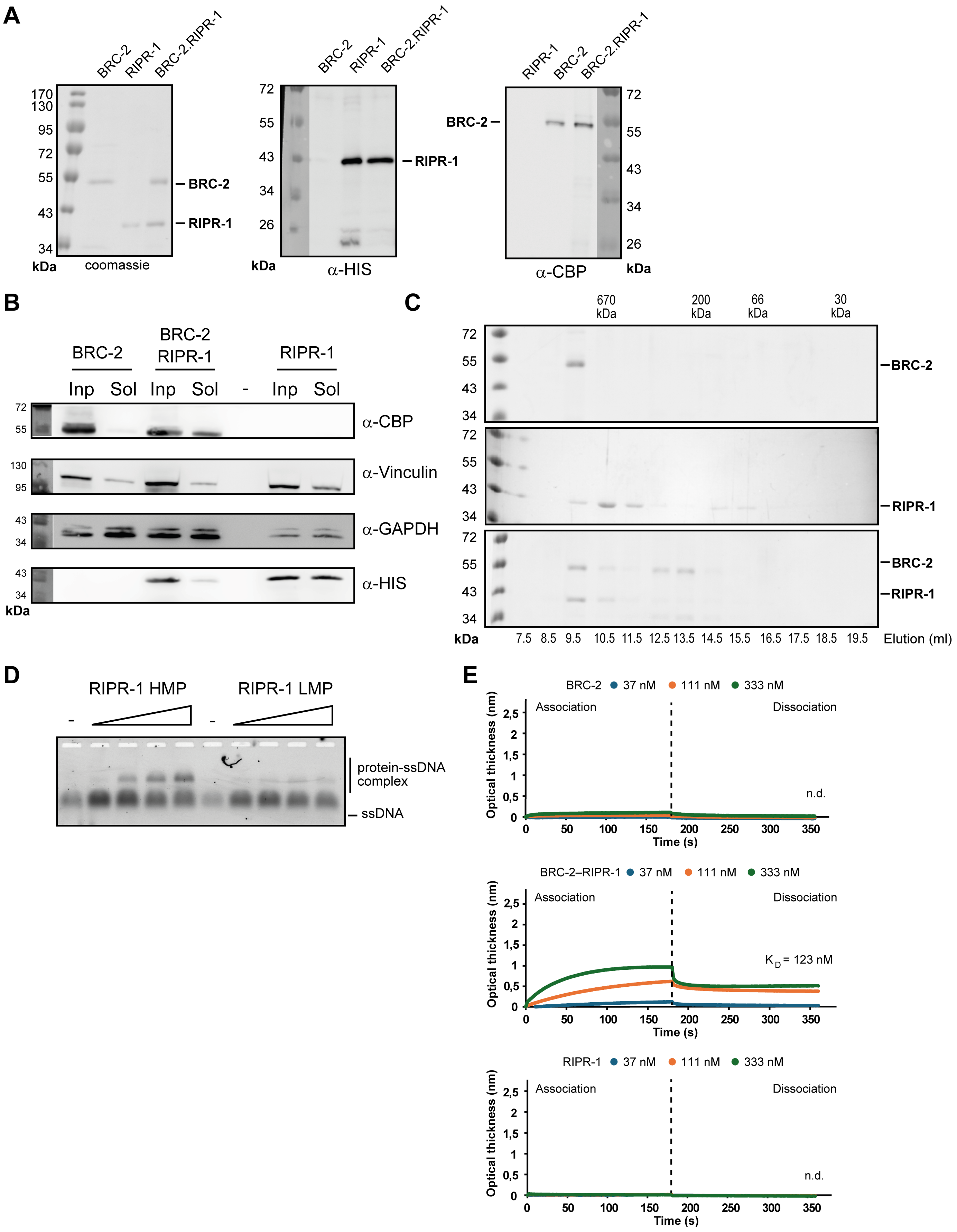
A
BRC-2
RIPR-1
BRC-2.RIPR-1
170
130
95
72
55
43
34
kDa
coomassie
BRC-2
RIPR-1
72
55
43
34
26
kDa
α-HIS
RIPR-1
BRC-2
α-CBP
kDa
B
BRC-2
BRC-2 RIPR-1
RIPR-1
Inp Sol Inp Sol - Inp Sol
α-CBP
α-Vinculin
α-GAPDH
α-HIS
130
95
kDa
C
670 kDa
200 kDa
66 kDa
30 kDa
7.5 8.5 9.5 10.5 11.5 12.5 13.5 14.5 15.5 16.5 17.5 18.5 19.5
Elution (ml)
D
RIPR-1 HMP
RIPR-1 LMP
protein-ssDNA complex
ssDNA
E
BRC-2 37 nM 111 nM 333 nM
BRC-2–RIPR-1 37 nM 111 nM 333 nM
RIPR-1 37 nM 111 nM 333 nM
Association
Dissociation
Optical thickness (nm)
Time (s)
n.d.
K D = 123 nM

**Supplementary Figure 7. RIPR-1 and BRC-2 directly associate *in vitro* and bind both ss- and dsDNA.**

**(A)** CBP-BRC-2 and HIS-RIPR-1 co-expression and co-purification shows direct interaction *in vitro*. **(B)** Western blot analysis on protein extracts expressing either CBP-BRC-2 or HIS-RIPR-1, as well as co-expressed BRC-2–RIPR-1 proteins in insect cells. Antibodies against tags and loading control proteins are indicated. Note that presence of RIPR-1 increases BRC-2 solubility. **(C)** Size-exclusion chromatography showing differential pools of RIPR-1 protein (high and low molecular weight) and complex formation with BRC-2 in a 1:1 ratio. **(D)** EMSA assay showing that the high molecular weight (HMP) but not lower molecular weight (LMP) RIPR-1 species are capable of binding ssDNA. **(E)** BLI assay showing that consistent with EMSA, only associated BRC-2–RIPR-1 can bind dsDNA.
