## Supplementary Table 3 for "BRC-2/BRCA2-RIPR-1 mediated constraints on homologous recombination execution are spatiotemporally regulated during meiosis"

**List of strains used in this study**

| **Strain** | **Genotype** | **Source** |
| --- | --- | --- |
| N2 | Wild type | CGC |
| NSV518 | *ripr-1(syb7129)* I*/hT2 [bli-4(e937) let-?(q782) qIs48]* (I;III) | This study |
| CA1423 | *mels8; spo-11::AID::3xFLAG ieSi38spo-11::AID::3XFLAG ieSi38* IV | CGC |
| CA1199 | *unc-119(ed3)* III*; ieSi38* IV*.* | CGC |
| NSV529 | *ripr-1(syb7129)* I*/hT2 [bli-4(e937) let-?(q782) qIs48]* (I;III); *spo-11::AID::3xFLAG ieSi38spo-11::AID::3XFLAG ieSi38* IV | This study |
| PHX6679 | *ripr-1[syb6679(ripr-1::AID::V5)]* I | This study |
| VC172 | *cep-1(gk138)* I | CGC |
| NSV532 | *ripr-1(syb7129) cep-1(gk138)* I*/hT2 [bli-4(e937) let-?(q782) qIs48]* (I;III) | This study |
| NSV517 | *ripr-1::AID::V5* I*; OLLAS::rpa-1* II*; ieSi38* IV | This study |
| NSV197 | \| *mre-11[iow45(mre-11::GFP::3XFLAG)]* V | Smolikove lab |
| NSV550 | *ripr-1(syb7129)* I*/hT2 [bli-4(e937) let-?(q782) qIs48]* (I;III); *mre-11::GFP::3XFLAG* V | This study |
| NSV172 | *rad-50[ddr32(rad-50::3XFLAG)]* V | Trivedi et al.; 2022 |
| NSV548 | *ripr-1(syb7129) I/hT2 [bli-4(e937) let-?(q782) qIs48] (I;III); rad-50::3XFLAG V* | This study |
| SSM540 | *dna-2::AID::FLAG* II*; ieSi38* IV | Hicks et al.; 2022 |
| SSM581 | *dna-2::AID::FLAG* II*; exo-1(tm1842)* III*; ieSi38* IV | Hicks et al.; 2022 |
| NSV721 | *ripr-1(syb7129)* I*/hT2 [bli-4(e937) let-?(q782) qIs48]* (I;III); *dna-2::AID::FLAG* II; *ieSi38* IV | This study |
| NSV724 | *ripr-1(syb7129)* I*/hT2 [bli-4(e937) let-?(q782) qIs48]* (I;III)*; dna-2::AID::FLAG* II*; exo-1(tm1842)* III*/[bli-4(e937) let-?(q782) qIs48]* (I;III)*; ieSi38* IV | This study |
| CA1199 | *unc-119(ed3)* III*; ieSi38* IV | CGC |
| NSV506 | *ripr-1::AID::V5* I*; ieSi38* IV | This study |
| PHX8244 | *ripr-1[syb8244(ripr-1::GFP)]* I | This study |
| DW104 | *brc-2(tm1086)* III*/hT2 [bli-4(e937) let-?(q782) qIs48]* (I;III) | CGC |
| NSV538 | *brc-2(syb7480)* III*/hT2 [bli-4(e937) let-?(q782) qIs48]* (I;III) | This study |
| NSV579 | *ripr-1(syb7129)* I*/hT2 [bli-4(e937) let-?(q782) qIs48]* (I;III);  *brc-2(syb7480)* III*/hT2 [bli-4(e937) let-?(q782) qIs48]* (I;III) | This study |
| NSV80 | *brc-2[ddr8(3xFLAG::brc-2)]* III | Trivedi et al.; 2022 |
| NSV114 | *brc-2[ddr15(HA::brc-2)]* III | This study |
| PHX9826 | *brc-2[syb9826(3XFLAG::AID::brc-2)]* III | This study |
| NSV666 | *3xFLAG::AID::brc-2* III*; ieSi38* IV | This study |
| NSV672 | *ripr-1::GFP* I*; 3XFLAG::AID::brc-2* III*; ieSi38* IV | This study |
| NSV709 | *brc-2[syb10557(3XFLAG::AID::brc-2^F35A^)]* III*/qC1* | This study |
| NSV716 | *ripr-1::GFP* I*; 3XFLAG::AID::brc-2^F35A^* III*/qC1* | This study |
| NSV97 | *cosa-1[ddr12(OLLAS::cosa-1)]* III | Janisiw et al.; 2018 |
| NSV510 | *ripr-1::AID::V5* I*; OLLAS::cosa-1* III*; ieSi38* IV | This study |
| NSV511 | *ripr-1::AID::V5* I*; HA::brc-2* III*; ieSi38* IV | This study |
| NSV512 | *ripr-1::AID::V5* I*; brc-2 (tm1086)/qC1* III*; ieSi38* IV | This study |
| NSV517 | *ripr-1::AID::V5* I*; OLLAS::rpa-1* II*; ieSi38* IV | This study |
| SSM476 | *rpa-1[iow92(OLLAS::rpa-1)]* II | Hefel et al.; 2021 |
| NSV531 | *ripr-1(syb7129)* I*/hT2 [bli-4(e937) let-?(q782) qIs48]* (I;III); *OLLAS::rpa-1* II | This study |
| NSV522 | *ripr-1::AID::V5* I*; cosa-1(tm3298)/qC1* III*; ieSi38* IV | This study |
| AV590 | *cosa-1(tm3298)/qC1* III | Yokoo et al.; 2012 |
| NSV624 | *cosa-1(tm3298) brc-2(syb7480)* III*/qC1* | This study |
| NSV523 | *ripr-1::AID::V5* I*; 3xFLAG::brc-2* III*; ieSi38* IV | This study |
| NSV549 | *ripr-1::AID::V5* I*; syp-2(ok307)* V*/nT1* (IV;V)*; ieSi38* IV*/nT1* (IV;V) | This study |
| QP2084 | *syp-3(ok758)/hT2 [bli-4(e937) let-?(q782) qIs48]* (I;III) | Yanowitz lab |
| NSV586 | *syp-3(ok758)* I*/hT2 [bli-4(e937) let-?(q782) qIs48]* (I;III); *brc-2(syb7480)* III/*hT2 [bli-4(e937) let-?(q782) qIs48]* (I;III) | This study |
| AV276 | *syp-2(ok307)* V*/nT1* (IV;V) | CGC |
| NSV129 | *msh-5[ddr22(GFP::msh-5)]* IV | Janisiw et al.; 2018 |
| NSV573 | *ripr-1(syb7129)* I*/hT2 [bli-4(e937) let-?(q782) qIs48]* (I;III); *GFP::msh-5* IV | This study |
| NSV688 | *rmh-1[syb9786(GFP::rmh-1)]* I | Engebrecht et al.; 2025 |
| NSV664 | *GFP::rmh-1* I*; brc-2(syb7480)/qC1* | This study |
| NSV527 | *ripr-1(syb7129)* I*/hT2 [bli-4(e937) let-?(q782) qIs48]* (I;III); *OLLAS::cosa-1* I/ *OLLAS::cosa-1 hT2 [bli-4(e937) let-?(q782) qIs48]* (I;III) | This study |
| NSV510 | *ripr-1::AID::V5* I; *OLLAS::cosa-1* III; *ieSi38* IV | This study |
| VC1873 | *rad-51(ok2218)* IV*/nT1 [qIs51]* (IV;V) | CGC |
| NSV720 | *ripr-1::AID::V5* I*; OLLAS::cosa-1* III*; rad-51(ok2218) ieSi38* IV*/ nT1* (IV;V) | This study |
| NSV545 | *ripr-1::AID::V5* I*; brc-2(syb7480)/qC1* III; *ieSi38* IV | This study |
| NSV528 | *ripr-1(syb7129)* I*/hT2 [bli-4(e937) let-?(q782) qIs48]* (I;III); *3XFLAG::brc-2* III/*hT2 [bli-4(e937) let-?(q782) qIs48]* (I;III) | This study |
| NSV584 | *ripr-1::GFP* I*/hT2 [bli-4(e937) let-?(q782) qIs48]* (I;III); *brc-2(syb7480)* III*/hT2 [bli-4(e937) let-?(q782) qIs48]* (I;III) | This study |
| PHX9833 | *ripr-1[syb9833(ripr-1::AID::3XHA)]* I | This study |
| NSV673 | *ripr-1::AID::3XHA* I*; 3XFLAG::brc-2* III*; ieSi38* IV | This study |
| PHX9826 | *brc-2[syb9826(3XFLAG::AID::brc-2)]* III | This study |
| NSV672 | *ripr-1::GFP* I*; 3XFLAG::AID::brc-2* III*; ieSi38* IV | This study |
| NSV511 | *ripr-1::AID::V5* I*; HA::brc-2* III | This study |
| NSV648 | *rad-51b[ske48(V5::rad-51b)]* IV | Neves et al.; 2025 |
| NSV715 | *ripr-1::GFP* I*; 3XFLAG::brc-2* III*; V5::rad-51b ieSi38* IV | This study |
