## Supplementary Table 4 for "BRC-2/BRCA2-RIPR-1 mediated constraints on homologous recombination execution are spatiotemporally regulated during meiosis"

**List of antibodies used in this study**

| **Antibody** | **Application (dilution)** | **Source** |
| --- | --- | --- |
| Monoclonal Mouse anti-HA | IF (1:600) | BioLegend (#901501) |
| Monoclonal Mouse anti-HA | WB (1:1000) | Cell Signaling (#2367) |
| Polyclonal Rabbit anti-HA | IF (1:1000) | Proteintech (81290-1-RR) |
| Monoclonal Rat anti-HA (HRP-conjugated) | WB (1:3000) | Roche (#12013819001) |
| Monoclonal Mouse anti-FLAG | IF (1:600) | Sigma (F1804) |
| Monoclonal Mouse anti-FLAG (HRP-conjugated) | WB (1:5000) | Sigma (A8592) |
| Polyclonal Rabbit anti-SYP-1 | IF (1:2000) | Janisiw et al.; 2020 |
| Polyclonal Rat anti-SYP-1 | IF (1:200) | Hicks et al.; 2022 |
| Polyclonal Rabbit anti-RAD-51 | IF (1:3000) | Das et al.; 2022 |
| Polyclonal Rat anti-RAD-51 | IF (1:500) | Blazickova et al.; 2025 |
| Polyclonal Rabbit anti-OLLAS | IF (1:1000) | Genscript (#A01658) |
| Monoclonal Mouse anti-GFP | IF (1:500) | Roche (#11814460001) |
| Polyclonal Rabbit anti-GFP 488-conjugated | IF (1:300) | Thermofisher (A-21311) |
| Polyclonal Guinea Pig anti-HTP-3 | IF (1:750) | Yumi Kim Lab |
| Polyclonal Guinea Pig anti-phosphorylated SUN-1^S8^ | IF (1:750) | Woglar et al.; 2013 |
| Polyclonal Rabbit anti V5 | IF (1:500) | Proteintech (14440-1-AP) |
| Monoclonal Mouse anti GAPDH | WB (1:5000) | Sigma (G8795) |
| Polyclonal Rabbit anti-histone H3 | WB (1:100,000) | AbCam (ab1791) |
| Polyclonal Chicken anti-GFP | WB (1:5000) | AbCam (ab13970) |
| Monoclonal Mouse anti-V5 | IF (1:500), WB (1:1000) | ThermoFisher (R960-25) |
| Monoclonal Mouse anti-Actin Clone C4 | WB (1:2000) | Santa Cruz (sc-47778) |
| Alexa Fluor anti-Rabbit 488 | IF (1:500) | ThermoFisher (#A-11034) |
| Alexa Fluor anti-Rabbit 594 | IF (1:750) | ThermoFisher (#A-11037) |
| Alexa Fluor anti-Mouse 488 | IF (1:500) | ThermoFisher (#A-11029) |
| Alexa Fluor anti- Mouse 594 | IF (1:750) | ThermoFisher (#A-11032) |
| Alexa Fluor anti-Rat 488 | IF (1:500) | ThermoFisher (#A-11006) |
| Alexa Fluor anti-Rat 555 | IF (1:750) | ThermoFisher (#A-21434) |
| Alexa Fluor anti-Guinea Pig 488 | IF (1:500) | ThermoFisher (#A-11073) |
| Alexa Fluor anti- Guinea Pig 594 | IF (1:750) | ThermoFisher (#A-11076) |
| Goat anti-Mouse HRP-conjugated | WB (1:15,000) | ThermoFisher (#31430) |
| Goat anti-Rabbit HRP-conjugated | WB (1:15,000) | ThermoFisher (#31460) |
| Goat anti-Guinea Pig HRP-conjugated | WB (1:15,000) | ThermoFisher (#A18769) |
| Goat anti-Rat HRP-conjugated | WB (1:15,000) | ThermoFisher (#31470) |
