## Supplementary Table 5 for "BRC-2/BRCA2-RIPR-1 mediated constraints on homologous recombination execution are spatiotemporally regulated during meiosis"

| **# of nuclei analysed for synapsis quantification** | **1** | **2** | **3** | **4** | **5** | **6** |
| --- | --- | --- | --- | --- | --- | --- |
| WT **(Supp. Fig. 1)** | 114 | 137 | 117 | 131 | 102 | 73 |
| *ripr-1(syb7129)* **(Supp. Fig. 1)** | 134 | 154 | 120 | 138 | 126 | 79 |

| **# of nuclei analysed for GFP::MSH-5 quantification** | **1** | **2** | **3** | **4** | **5** |
| --- | --- | --- | --- | --- | --- |
| *GFP::msh-5* **(Fig. 5)** | 86 | 76 | 77 | 43 | 61 |
| *ripr-1(syb7129); GFP::msh-5* **(Fig. 5)** | 227 | 235 | 217 | 151 | 104 |

| **# of nuclei analysed for GFP::RMH-1 quantification** | **1** | **2** | **3** | **4** | **5** |
| --- | --- | --- | --- | --- | --- |
| *GFP::rmh-1* **(Fig. 5)** | 191 | 144 | 114 | 87 | 70 |
| *GFP::rmh-1; brc-2(syb7480)* **(Fig. 5)** | 238 | 207 | 189 | 137 | 96 |

| **# of Diakinesis nuclei analysed for quantification of DAPI bodies** | **n** |
| --- | --- |
| WT **(Fig. 1)** | 49 |
| *ripr-1(syb7129)* **(Fig. 1)** | 51 |
| *spo-11::AID ieSi38*  (-auxin) **(Fig. 2)** | 39 |
| *ripr-1(syb7129); spo-11::AID ieSi38*(-auxin) **(Fig. 2)** | 53 |
| *spo-11::AID ieSi38*  (+auxin) **(Fig. 2)** | 30 |
| *ripr-1(syb7129); spo-11::AID ieSi38* (+auxin) **(Fig. 2)** | 41 |

| **# of nuclei analysed for COSA-1 quantification** | **# of nuclei** |
| --- | --- |
| *OLLAS::cosa-1* **(Fig. 5)** | 136 |
| *ripr-1(syb7129); OLLAS::cosa-1* **(Fig. 5)** | 168 |
| *OLLAS::cosa-1 brc-2(syb7480)* **(Fig. 5)** | 160 |
| *ripr-1::AID::V5; OLLAS::cosa-1 ieSi38*  (-auxin) **(Fig. 5)** | 116 |
| *ripr-1::AID::V5; OLLAS::cosa-1 ieSi38*  (+auxin) **(Fig. 5)** | 152 |
| *ripr-1::AID::V5; OLLAS::cosa-1 rad-51(ok2218) ieSi38*  (-auxin) **(Fig. 5)** | 257 |
| *ripr-1::AID::V5; OLLAS::cosa-1 rad-51(ok2218) ieSi38*  (+auxin) **(Fig. 5)** | 240 |

| **# of nuclei analysed for RAD-51 foci quantification** | **1** | **2** | **3** | **4** | **5** | **6** | **7** |
| --- | --- | --- | --- | --- | --- | --- | --- |
| WT **(Fig. 1)** | 259 | 356 | 300 | 246 | 237 | 199 | 143 |
| *ripr-1(syb7129)* **(Fig. 1)** | 266 | 323 | 301 | 273 | 216 | 175 | 120 |
| *spo-11::AID ieSi38* (-auxin) **(Fig. 2)** | 160 | 205 | 157 | 159 | 141 | 115 | 94 |
| *ripr-1(syb7129); spo-11::AID ieSi38*  (-auxin) **(Fig. 2)** | 241 | 350 | 283 | 245 | 210 | 163 | 120 |
| *spo-11::AID ieSi38* (+auxin) **(Fig. 2)** | 352 | 370 | 332 | 281 | 223 | 188 | 138 |
| *ripr-1(syb7129); spo-11::AID ieSi38*  (+auxin) **(Fig. 2)** | 141 | 158 | 146 | 126 | 110 | 85 | 66 |
| *spo-11::AID ieSi38* (-auxin) **(Fig. 3)** | - | - | - | - | - | - | 96 |
| *spo-11::AID ieSi38* (+auxin) **(Fig. 3)** | - | - | - | - | - | - | 69 |
| *ripr-1(syb7129); spo-11::AID ieSi38*  (-auxin) **(Fig. 3)** | - | - | - | - | - | - | 50 |
| *ripr-1(syb7129); spo-11::AID ieSi38*  (+auxin) **(Fig. 3)** | - | - | - | - | - | - | 60 |
| *dna-2::AID ieSi38*  (-auxin) **(Fig. 3)** | - | - | - | - | - | - | 76 |
| *dna-2::AID; exo-1(tm1842); ieSi38*  (-auxin) **(Fig. 3)** | - | - | - | - | - | - | 82 |
| *ripr-1(syb7129); dna-2::AID ieSi38*  (-auxin) **(Fig. 3)** | - | - | - | - | - | - | 70 |
| *ripr-1(syb7129); dna-2::AID;*  *exo-1(tm1842); ieSi38*  (-auxin) **(Fig. 3)** | - | - | - | - | - | - | 102 |
| *dna-2::AID ieSi38*  (+auxin) **(Fig. 3)** | - | - | - | - | - | - | 85 |
| *dna-2::AID; exo-1(tm1842); ieSi38*  (+auxin) **(Fig. 3)** | - | - | - | - | - | - | 74 |
| *ripr-1(syb7129); dna-2::AID ieSi38*  (+auxin) **(Fig. 3)** | - | - | - | - | - | - | 55 |
| *ripr-1(syb7129); dna-2::AID;*  *exo-1(tm1842); ieSi38*  (-auxin) **(Fig. 3)** | - | - | - | - | - | - | 107 |
| WT **(Fig. 4)** | 153 | 239 | 222 | 183 | 160 | 139 | 98 |
| *ripr-1(syb7129)* **(Fig. 4)** | 187 | 188 | 203 | 191 | 165 | 98 | 69 |
| *brc-2(syb7480)* **(Fig. 4)** | 178 | 212 | 188 | 167 | 157 | 100 | 49 |
| *ripr-1(syb7129); brc-2(syb7480)*  **(Fig. 4)** | 162 | 203 | 180 | 144 | 143 | 110 | 55 |
| *ripr-1::AID::V5; ieSi38*  (-auxin) **(Fig. 4)** | 160 | 248 | 201 | 143 | 125 | 118 | 89 |
| *ripr-1::AID::V5; cosa-1(tm3298); ieSi38*  (-auxin) **(Fig. 4)** | 173 | 211 | 179 | 157 | 127 | 107 | 96 |
| *ripr-1::AID::V5; ieSi38*  (+auxin) **(Fig. 4)** | 164 | 242 | 187 | 178 | 153 | 95 | 62 |
| *ripr-1::AID::V5; cosa-1(tm3298); ieSi38*  (+auxin) **(Fig. 4)** | 103 | 129 | 115 | 104 | 92 | 61 | 42 |
| WT **(Fig. 4)** | 98 | 163 | 109 | 99 | 85 | 77 | 64 |
| *brc-2(syb7480)* **(Fig. 4)** | 187 | 212 | 183 | 188 | 169 | 104 | 67 |
| *syp-3(ok758)* **(Fig. 4)** | 158 | 183 | 138 | 122 | 103 | 92 | 97 |
| *syp-3(ok758);* *brc-2(syb7480)* **(Fig. 4)** | 122 | 131 | 106 | 107 | 88 | 59 | 33 |
